# Studies of vegetation and land-use history on Bodmin Moor: pollen investigations from Leskernick

**DOI:** 10.1101/2024.01.05.574344

**Authors:** Martyn Waller

## Abstract

Details of pollen investigations that accompanied archaeological excavations (of a Late Neolithic/Early Bronze ritual complex and a Middle to Late Bronze Age settlement) at Leskernick Hill Bodmin Moor, undertaken between 1995 and 1999, are presented. Samples were collected both from a nearby ‘off-site’ location and from a variety of ‘on-site’ contexts. The off-site peat profile contains a major hiatus. The earliest assemblage corresponds to the Late Neolithic/Early Bronze Age and indicates a largely open landscape in which both pasture and arable agriculture were practiced. Sedimentation recommenced during the early Medieval period and the pollen record shows the varying degrees to which grazing and arable agriculture were practiced up until the recent past. These phases do not accord well with historic evidence from the wider area, though other pollen diagrams from the region also record a wetter phase found at Leskernick (and dated here to *c*. 1400-1600 AD). Evidence obtained during the archaeological excavations suggests the ritual complex was abandoned early and the pollen records indicate this was accompanied by a period of low grazing pressure. Samples from the interiors of huts/houses dated to the Middle Bronze Age are dominated by Poaceae pollen suggesting relatively intense human activity. The assemblages from beneath the boundary walls around the settlement probably cover a considerable time period, in part as a result of rebuilding and the reuse of stone. With high *Calluna vulgaris* values they appear largely to reflect the stony nature of the environment in which the settlement was established.

**Preamble:** This pre-print details the results of pollen investigations that accompanied archaeological excavations at Leskernick Hill, Bodmin Moor, undertaken between 1995 and 1999. A draft manuscript was prepared for publication in 2005, but the monograph in which it was to be included outlining the excavations, has not been forthcoming. Details of the archaeological/historic features present at Leskernick can be found in Johnson and Rose (1994) and Herring *et al*. (2008), while Bender *et al*. (2007) provides information on the work that these investigations formed part of. The 2005 manuscript is reproduced here with a new introduction and revisions incorporating information from the later publications.

## Introduction

Bodmin Moor comprises an area of approximately 200 km^2^ of granitic upland in east Cornwall south-west England, the high tors of which attain >300m OD. As a result of its westerly position Bodmin Moor experiences a mild and wet climate, with precipitation at about 1800 mm annually. The higher areas are covered in unimproved acid grassland, typically developed over stagnohumic gley soils were a thin humic topsoil overlies a sandy loam of variable permeability (Cranfield University 2023). There are numerous rocky outcrops (tors and boulder debris fields), scattered heathlands and, in the associated shallow valleys, mires. Pastoralism, especially sheep, is the main agricultural activity, though since 1945 the area has seen changes in grazing regime, agricultural intensification, reservoir construction and some afforestation (Jones and Essex 1999).

Johnson and Rose (1994) detail the wealth of archaeological remains found on Bodmin Moor. These include prehistoric defended enclosures and monuments. While some of former date from the Neolithic, activity is thought to have peaked during the Bronze Age. The monuments from this period include cairns (over 300 are present), the larger of which are sited in prominent positions, while the small (<5m diameter) occur around the ritual monuments. The latter include stone circles (16) and stone rows (7). There are also a large number of presumed Bronze Age settlements, some consisting of unenclosed huts (the term preferred by Johnson and Rose over that of ‘house’ as it does not imply habitation) and others with huts both around and within enclosures. Such enclosures can be difficult to distinguish from prehistoric field boundaries (e.g. boundaries that define areas of controlled grazing or arable land). Bodmin Moor also has an abundance of historic remains; 37 deserted medieval settlements and 227 field systems some of which absorbed their prehistoric equivalents. Place name evidence suggests the early practice of transhumance. The region appears to have been permanently settled after 1100 AD with most settlements recorded by the 14^th^ century. This period, during which arable activity was practised, represents a medieval ‘high tide’ as desertions and settlement shrinkage occurred from the 15^th^ century onwards. A final phase of post-medieval expansion is particularly associated with industrial activity; mineral (notably tin) and china clay workings and processing sites and granite quarrying (Herring *et al*. 2008).

The site of the investigations reported here, Leskernick Hill, rises to elevation of 300 m OD. At the base of the hill is a Late Neolithic/Early Bronze ritual complex consisting of two stone circles and a stone row (Figure 1). A Middle to Late Bronze Age settlement occurs on the south facing slope. This comprises *c*. 50 huts and enclosures/field walls amongst an extensive spread of boulder debris/‘clitter’ (Bender *et al*. 2007, Hamilton *et al*. 2008). The boundary systems are typical for the Bronze Age, that is they are “curvilinear and accreted, having developed organically from one or more foci” (Johnson and Rose 1994 p59).

**Figure 1.**
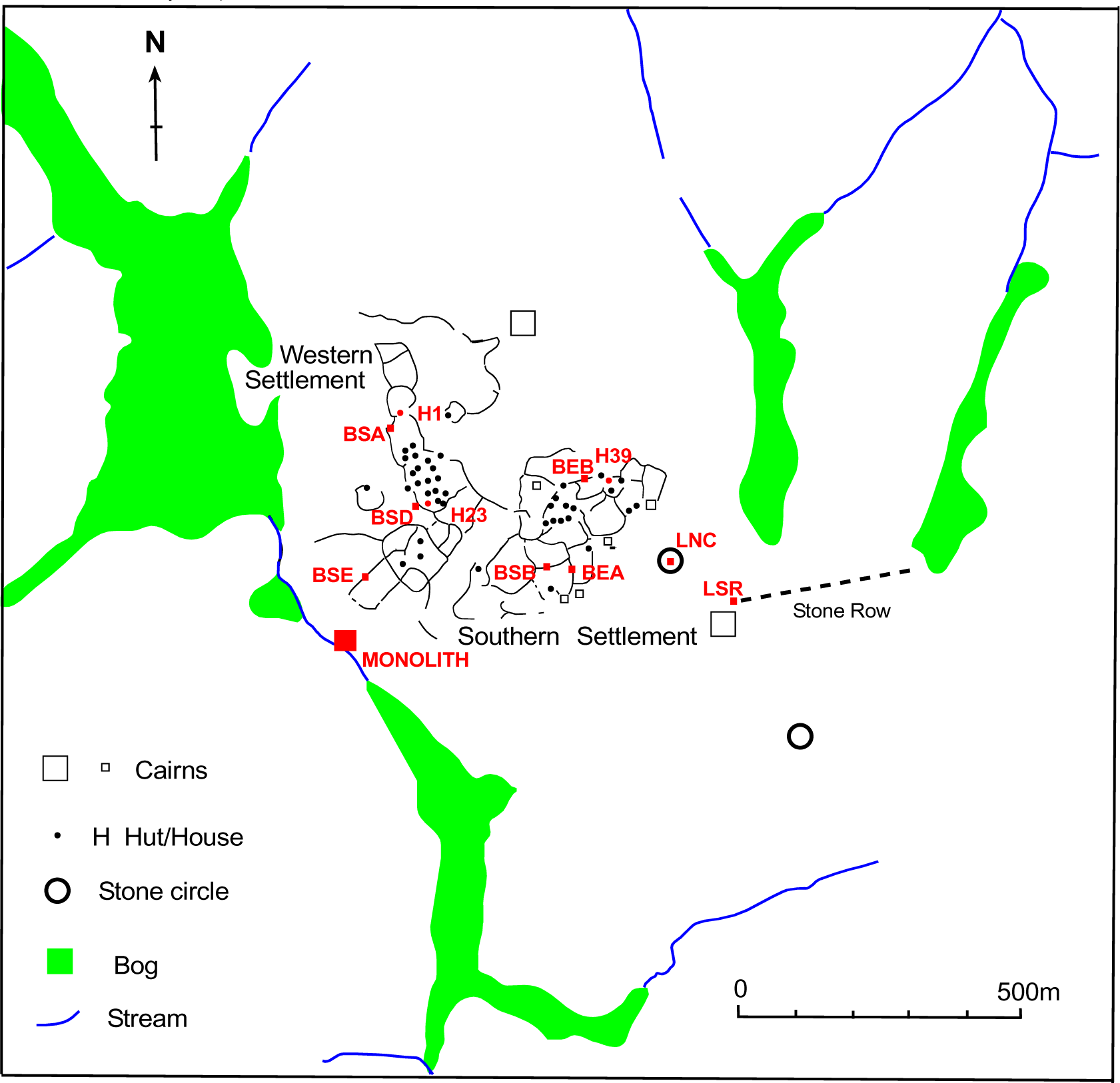
Prehistoric Leskernick: showing the settlements, cairns, stones row, circles and the location of the pollen samples (in red, abbreviations follow the text). After Johnson and Rose (1994). The large ‘cairn’ situated near the western end of the Stone Row is now considered likely to be the result of mining activity (Bender *et al*. 2007).

The lower slopes of a hillside to the south-west of the prehistoric remains were subject to arable agriculture in the medieval period (Figure 2) with oats, roots, brassicas among the likely crops (Herring *et al*. 2008). Herring *et al*. (2008) also detail the extensive post Medieval remains found at Leskernick. These include a settlement (‘Leskernick’) and associated ‘pasture’ field system (1808-*c*.1840) and numerous mineral workings. Associated with the surrounding valley bottoms are ‘streamworks’ (tin workings characterised by extensive cuttings and drainage gullies containing dumps of waste), along spreads of both eluvial (material detached and moved downslope by periglacial activity during the Quaternary) and alluvial deposits, and a network of leats. The deposits at the top of the neighbouring Codda Tor were subject to turf cutting (Herring *et al*. 2008).

**Figure 2.**
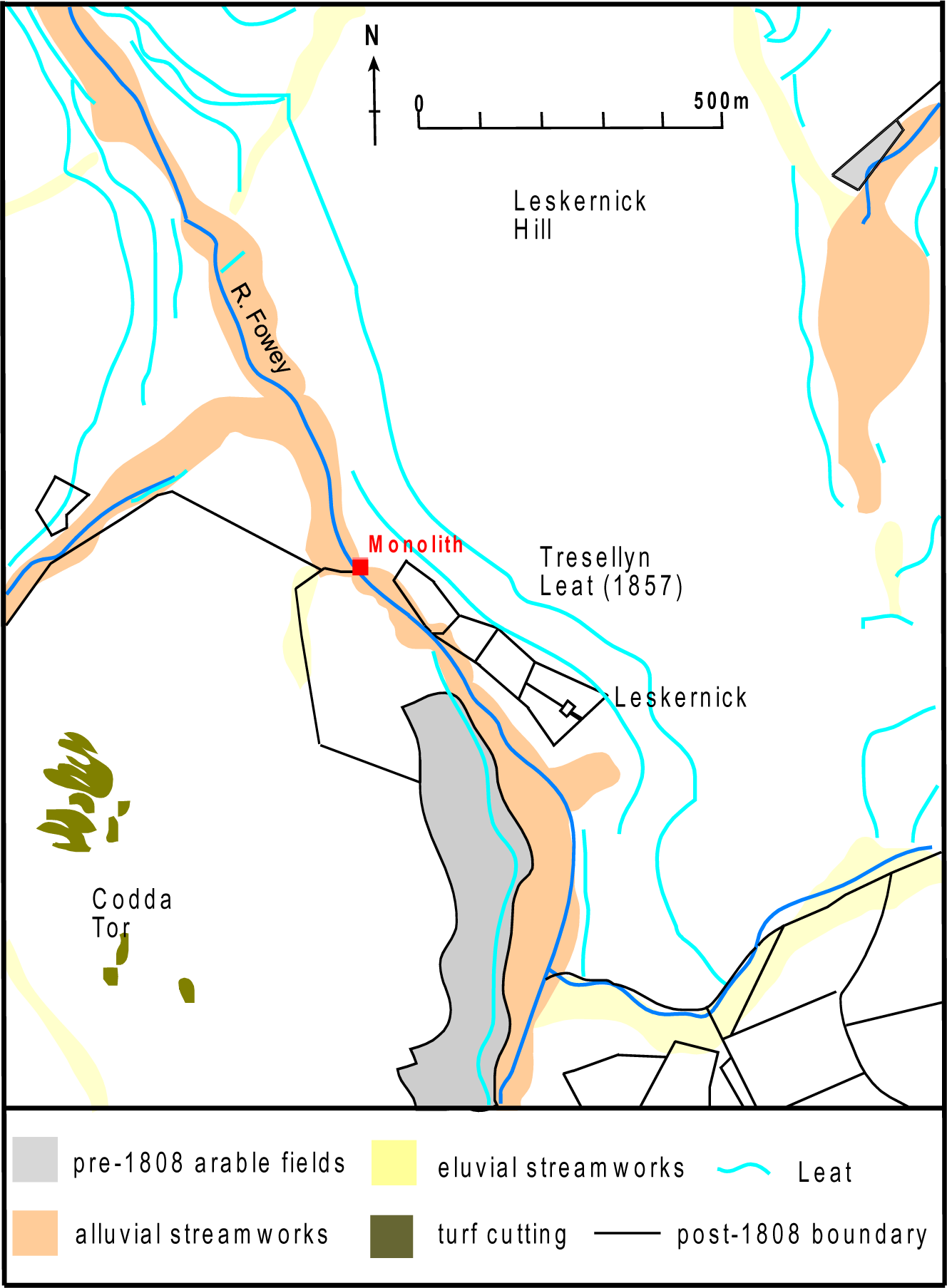
Medieval Leskernick: showing the location of the pollen monolith. After Herring *et al*. (2008).

The aims of the pollen investigations described here were 1) to obtain a vegetation/land-use record from the vicinity of the archaeological remains at Leskernick that spanned the period of pre-historic activity, utilising nearby (‘off-site’) organic deposits and 2) in conjunction with the archaeological excavations to reconstruct the environment (alongside other environmental analysis, phosphates and charcoal) from archaeological contexts and other sealed surfaces (e.g. below archaeological remains).

Plant nomenclature follows Stace (1997).

### Previous pollen investigations of vegetation and land-use history from Bodmin Moor

Research into the past vegetation of Bodmin Moor has focused on a number of themes. These include the presence and nature of woodland in the early Holocene, changes in woodland cover during the Neolithic and the vegetation and land-use associated with the occupation of the area during the later prehistoric and historic periods. Pollen based reconstructions at the landscape scale have been hindered here and the Cornish uplands in general by the lack of detailed radiocarbon dated Holocene pollen records. The reasons, detailed by Caseldine (1980; 1983), include both the absence of deep glacial basins and disturbance, particularly peat cutting and streaming associated with the tin industry.

Early work from Bodmin Moor (Conolly *et al*. 1950; Brown 1977) focused on Late Glacial and early Holocene deposits located at Parsons Park, Hawk Tor, Stannon Marsh and Dozmary Pool (Figure 3). The work of Brown (1977) has been particularly influential, with his interpretation of low Holocene tree and shrub pollen values as indicating the absence of woodland cover over the higher areas of the moor, widely accepted. Changes in peat stratigraphy in the upper parts of the profiles were attributed to climatic deterioration. Unfortunately, the sequences at Parsons Park and Hawks Tor (both at *c*. 230 m OD) were, above breaks in sedimentation which occurred during early Holocene, not independently dated, the date of 5530-5230 cal. yrs BC from Dozmary Pool (at 265 m OD) being the youngest obtained. Only the latter site has subsequently been re-examined (Simmons *et al*. 1987), a study which suggests that the stratigraphic integrity of the later deposits here can be questioned. In addition, as Geary *et al*. (2000a) have commented, the location of the Dozmary Pool sequence (on a mire adjacent to the pool), some distance away from any potentially wooded areas and with the local mire vegetation inevitably well represented in the pollen rain, may account for the low percentages of tree and shrub pollen.

**Figure 3.**
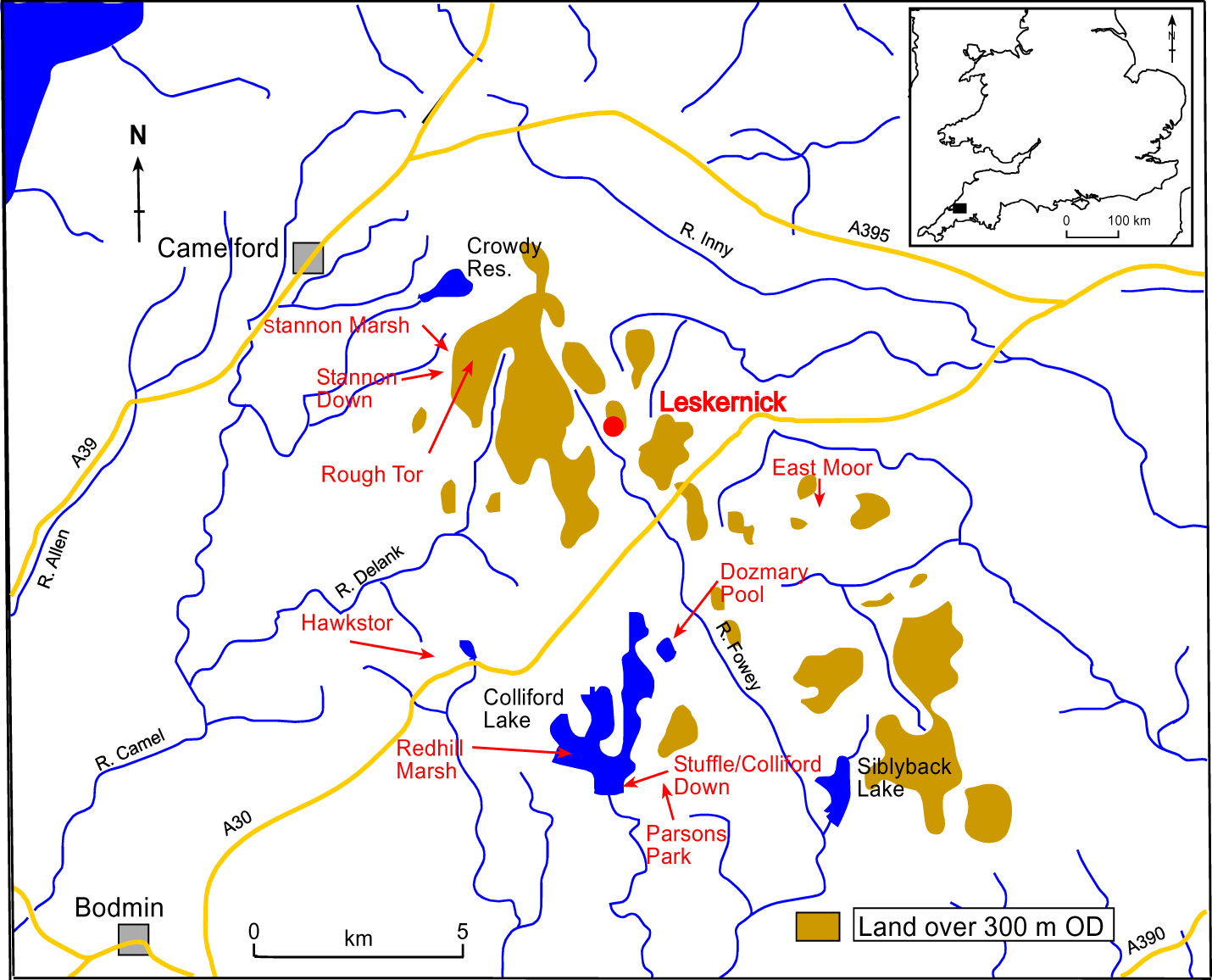
Bodmin Moor: showing sites of previous pollen studies and the location of Leskernick. Inset the location of Bodmin Moor (black rectangle).

More recently, Geary *et al*. (2000a, b) have published a number of pollen sequences from wetland areas in the vicinity of Rough Tor (at *c*. 300 m OD) and East Moor (at *c*. 280 m OD) which date from *c*. 8200-7000 cal. yrs BC onwards. High tree (with maximum *Quercus* values of over 20% TLP) and shrub (with maximum *Corylus avellana*-type values of over 90% TLP) pollen percentages provide evidence of dense woodland cover on the adjacent hillsides, which Geary *et al*. (2000a) suggest probably extended up to the summits prior to *c*. 5500 cal. yrs BC (Geary *et al*. 2000a). The valley bottoms also appear to have been wooded after *c*. 4750 cal. yrs BC when values for *Alnus glutinosa* rise suggesting increased soil moisture, possibly as a consequence of human activity during the Mesolithic. Increases in Poaceae and *Potentilla*-type in Rough Tor South sequence, indicative of small-scale disturbance, could also be linked to such activity (Geary *et al*. 2000a). Unfortunately while convincingly indicating the presence of woodland, there are problems with the chronology for the basal assemblages at both Rough Tor South (with an age inversion) and the East Moor monolith, were only the top of the EMM1 sequence is dated. Pollen assemblages at Redhill Marsh (Collingford, at *c*. 240 m OD) also indicate the local establishment of mixed deciduous woodland (with *Betula*, *Quercus* and *Corylus* the main taxa) by 7960-7530 cal. yrs BC, though here the later assemblages are undated and breaks in sedimentation may have occurred (Walker and Austin 1985).

In Geary *et al*. (2000b) the first clear evidence of human activity is taken to be the onset of the consistent presence of *Plantago lanceolata* pollen (indicating the creation and maintenance open areas), which occurs before *c*. 3500 cal. yrs BC in the Rough Tor South sequence. Pastoral activity is indicated, though the grazing pressure envisaged is light and possibly intermittent. After initial declines, *Corylus avallena*-type pollen percentages recover in both the Rough Tor South and East Moor monolith sequences. Dated to *c*. 2700 cal. yrs BC at East Moor, this phase of regeneration and therefore reduced grazing pressure, is seen by Geary *et al*. (2000b) as being the result of the focus of activity moving from the moor to the fringing valleys and as part of a wider shift in land use away from the uplands in the late Neolithic.

The Bronze Age is probably represented at five of the Geary *et al*. (2000a; 2000b) sites, though the temporal resolution of this period is consistently poor. Three of the sites indicate woodland clearance, with this activity directly dated at Rough Tor South (1670-1430 cal. yrs BC) and East Moor (1230-900 cal. yrs BC, Tresellern Marsh A). Removal of both the hazel on the slopes and the alder in valley bottoms is evident. The other sites suggest a more complex, presumably local variable, pattern of woodland exploitation. At Tresellern Marsh B (East Moor) major clearance appears to have occurred prior to 2880-2470 cal. yrs BC, while the Rough Tor North monolith C sequence indicates that areas of woodland persisted throughout the period *c*. 4400 to 600 cal. yrs BC (the RTNC1 assemblage). The herbaceous pollen types recorded from both the Rough Tor and East Moor sites during the Bronze Age (notably *Plantago lanceolata*, *Galium*-type, *Ranunculus acris*-type and *Potentilla*-type) are indicative of a predominantly pastoral landscape during the development of the adjacent field systems, though Cerealia-type pollen was consistently recorded from *c*. 2000 cal. yrs BC at East Moor (Tresellern Marsh A), suggesting the cultivation of cereals nearby (Geary *et al*. 2000b).

The evidence from peat-profiles of land-use during the Neolithic and Bronze Age can be supplemented by palynological data from archaeological contexts and soil profiles. Mercer and Dimbleby (1978) report investigations from a number of contexts associated with the Stannon Down Hut Circle Settlement. A layer previously (Mercer 1970) described as a cultivated soil (with quernstones excavated from the settlement), contained no cereal pollen and few arable weeds, though as noted by Dimbleby this does not necessarily negate the original interpretation, as conditions are unlikely to have been conducive to pollen preservation while the soil was being actively cultivated. A sample from a gutter fill contained a high proportion of *Quercus* and *Corylus* pollen and was thought to have been derived from the collapse of the gutter walls. The peaty layer above the ‘cultivated’ soil was also investigated, which Dimbleby suggests started to form sometime after cultivation ceased, cautioning against assuming that peat formation and abandonment occurred as a result of climate deterioration. Brisbane and Clew (1979) present a number of pollen sequences associated with a cairn and field system at East Moor. Samples were collected from a soil profile beneath a large flat stone at the base of a presumed Bronze Age cairn. High basal *Quercus* percentages were interpreted as representing local oak woodland, with the upper part of the profile suggesting both woodland clearance and podsolization occurred prior to cairn construction. In a nearby soil pit, high Ericaceae and Poaceae pollen values coincide with a peaty humus layer found beneath the modern turf, with an immediately preceding rise in *Pteridium* spores thought to represent a decline in grazing pressure prior to ‘peat’ formation. Brisbane and Clew (1979) therefore suggest, as alternative to the traditional view that agricultural abandonment occurred as a result of climatic deterioration and peat formation in the late Bronze Age, that progressive soil improvishment produced first a decline in grazing and eventually peat formation. Pollen data, from similar contexts, are also available from the Colliford Down area (Maltby and Caseldine 1982; Caseldine 1983; Griffith *et al*. 1984). The investigation of profiles beneath two barrows (CRII and CRIV, at 240-250 m OD) provides evidence of local variations in soil development during prehistory. The sequence beneath CRII, on a west-south-west facing slope, suggests that a brown earth, or possibly humic podzol, soil survived woodland clearance and grassland development, prior to the construction of the barrow (dated to 2145-1770 cal. yrs BC). On the opposite, east-north-east facing slope, an iron pan stagnopodzol developed (with grassland succeeded by heathland) prior to the construction of CRIV (dated to 2035-1620 cal. yrs BC). On the latter slope soil profiles not buried beneath archaeological features appear similar suggesting they have undergone little subsequent change. High basal *Corylus avellana*-type pollen frequencies from such soil profiles (on both slopes) were used to suggest hazel shrub may have increased in the area sometime after the barrows were constructed.

The Rough Tor pollen diagrams provide evidence for settlement abandonment and woodland regeneration on Bodmin Moor in later prehistory (Geary *et al*. 2000b). High tree pollen frequencies occur at the base of Rough Tor North monolith A (though they were not directly dated) and tree and shrub pollen frequencies rise at the top of the RTNC1 assemblage. The Rough Tor South sequence also indicates a phase of woodland regeneration though this appears to post-date the Roman period (after 250-540 cal. yrs AD). Geary *et al*. (2000b) caution, in the absence of local evidence, against adopting a model of abandonment occurring either as a result of climate change or soil deterioration. They argue the late Bronze Age/Iron Age could, in terms of vegetation cover, have been a period of continuity, with the low intensity grazing they envisage maintaining the open grasslands created earlier in the Bronze Age. Open conditions are certainly indicated during the Romano-British period at Rough Tor (e.g. Rough Tor monoliths A and C) and East Moor (e.g. Watery Marsh, Tresellern Marsh A). The herb-rich pollen assemblages are said to be indicative of the presence of meadows which were cut for hay and grazed in winter (Geary *et al*. 2000b).

The diagrams from Rough Tor and East Moor also contain information on the post-Roman development of the landscape. In the Rough Tor North monolith C sequence a herb-rich assemblage is replaced at 1160-1300 cal. yrs AD by one indicative of more intensive land-use with the occurrence of Cerealia-type pollen indicating arable agriculture occurring nearby (Geary *et al*. 1997). At East Moor (Tresellern Marsh A), slight increases in tree and shrub pollen are seen as representing a short-lived reduction, before a re-expansion, in agricultural activity, with Cerealia-type pollen again recorded (both phases are dated approximately to 890-1220 cal. yrs AD). A more detailed peat pollen profile is available from Stuffle near Colliford (at *c*. 260 m OD), from where soil pollen analyses were also undertaken (Walker in Austin *et al*. 1989). The base of Stuffle sequence indicates open grassland with limited tree cover. This is followed by a phase with higher *Calluna* pollen values (taken to indicate drier conditions) with Cerealia pollen grains present, the mid-point of which is dated to 1040-1270 cal. yrs AD. A subsequent period when mixed agriculture, with higher Cerealia pollen values, was clearly being locally practised is dated to 1280-1460 cal. yrs AD, with these high Cerealia pollen values recorded until 1400-1640 cal. yrs AD. The associated soil pollen assemblages appear to confirm an early phase of cultivation and suggest the period when arable agriculture is so prominent in the Stuffle profile, it was also at its most extensive. Interestingly, a similar phase to the latter, which has been recorded from a small bog within a field system at Okehampton Park (*c*. 340 m OD) on the northern edge of Dartmoor (Austin *et al*. 1980), commenced 1180-1410 cal. yrs AD. The uppermost assemblages in the Stuffle profile are thought to represent a period when the nearby farmsteads were deserted and a purely pastoral economy returned, followed by phase of more intensive activity, which may be related the use of the Colliford area during the 19^th^ century for cattle grazing (Walker in Austin *et al*. 1989). These together these investigations support the concept of a ‘high tide’ of activity on Bodmin Moor and possibly other uplands areas in south-western England, in the late Medieval period (Geary *et al*. 1997).

## Methods

Standard laboratory techniques were used to prepare samples of a known volume (generally 1 cm^3^) for pollen analysis (Moore *et al*. 1991), with *Lycopodium* tablets added to enable the calculation of pollen concentrations (Stockmarr 1971). Each sample was counted until a sum of not less than 500 total land pollen (TLP) grains had been exceeded in the off-site samples and not less than 300 TLP in the on-site samples (with the actual sums attained shown on the pollen diagrams). Pollen and spores were identified with the aid of reference material held at Kingston University. Nomenclature for pollen taxa follows Bennett (1994). Differentiation of the Poaceae group follows Andersen (1979a). Grains with annulus diameters >10 µm are equivalent to ‘Cerealia-type’ used by other Bodmin Moor palynologists. The preservation status of each grain was recorded following Cushing (1967) with proportions well preserved, determinable but deteriorated, and indeterminable presented here. Microscopic charcoal frequency was assessed using the point counting method of Clark (1982).

The pollen diagrams have been drawn using the computer programs TILIA and TILIA*GRAPH (Grimm 1993). A percentage of total land pollen sum (% TLP) has been used in their construction with aquatics expressed as a percentage of total pollen (% TP), pteridophytes as % TLP+pteridophytes and *Sphagnum* as % TLP+*Sphagnum*. Zonation of the off-site monolith and sequence from the Northern Stone Circle were undertaken using stratigraphically constrained cluster analysis performed by the program CONISS (Grimm 1987).

The pollen datasets from Leskernick have been summarised with the aid of an ordination technique. Detrended correspondence analysis (DCA) was performed on the on-site and off-site samples separately, using the CANOCO 4 program (ter Braak and Šmilauer 1998).

Five samples from the off-site location were submitted to Beta Analytic for radiocarbon dating by Accelerator Mass Spectrometry (AMS). All the calibrated date ranges quoted in the text (at the two sigma age range) were calculated using the IntCal20 dataset in the program CALIB v. 8.1.0 (see Stuiver and Reimer 1993).

### Off-site investigations

#### Introduction

The off-site investigations were undertaken from a site (NGR SX17917986) in the bottom of the valley (at *c*. 260 m OD) located to the west of the Western Settlement (Figures 1 and 2). The peat deposits on the valley floor are thin, though an exposed stream section revealed an organic sequence 110 cm in depth. Material was collected from the cleaned exposure using a monolith tin (10×10×100 cm), which was stored at 3^0^C prior to sub-sampling for pollen and the extraction of material for radiocarbon dating. The lithology at this site is detailed in Table 1.

**Table 1.**
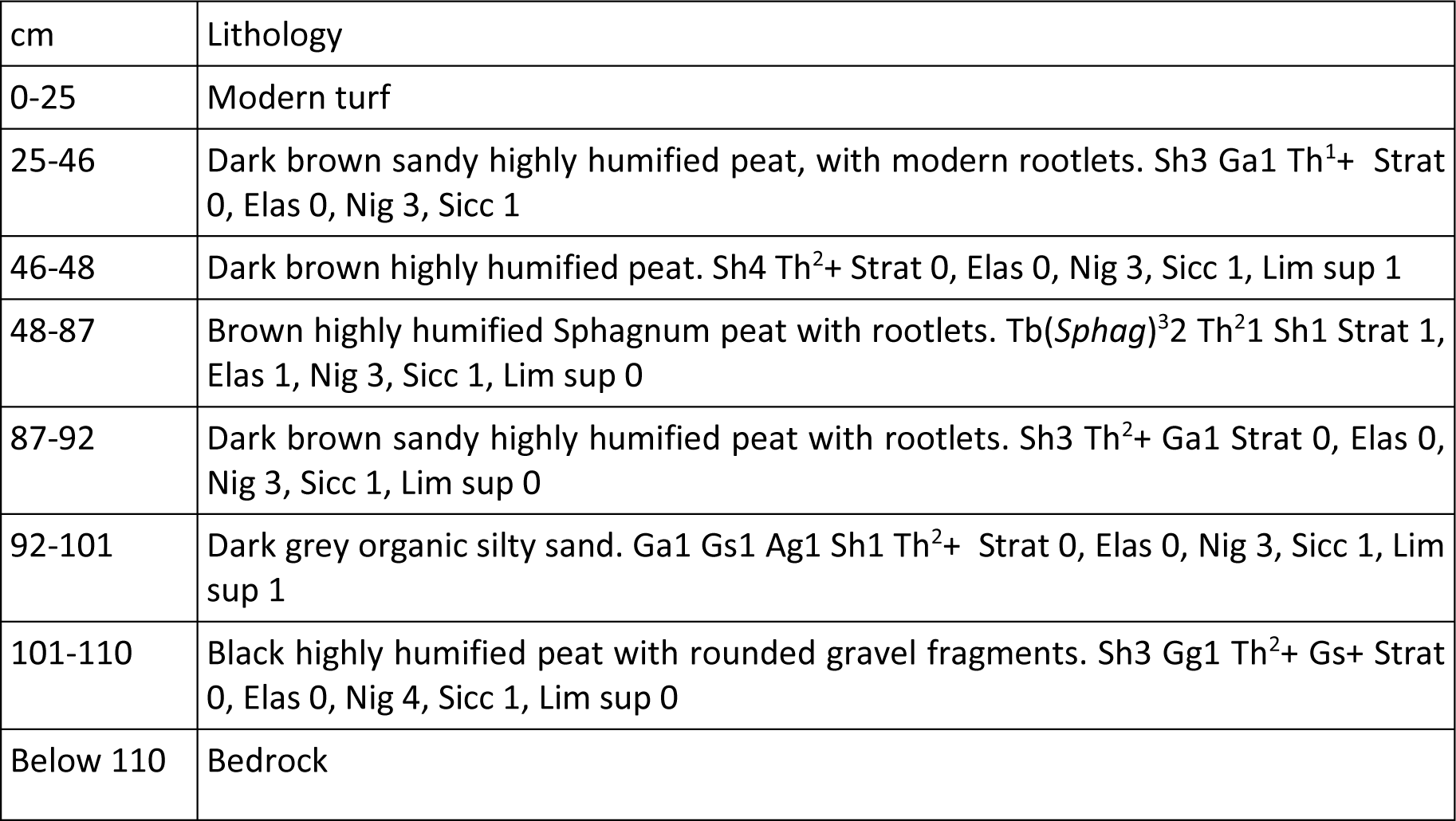
Lithology of the pollen monolith (notation follows Troels-Smith 1955)

The off-site pollen diagram (Figures 4 and 5) has been divided into five local pollen assemblage zones prefixed LESKM.

**Figure 4:**
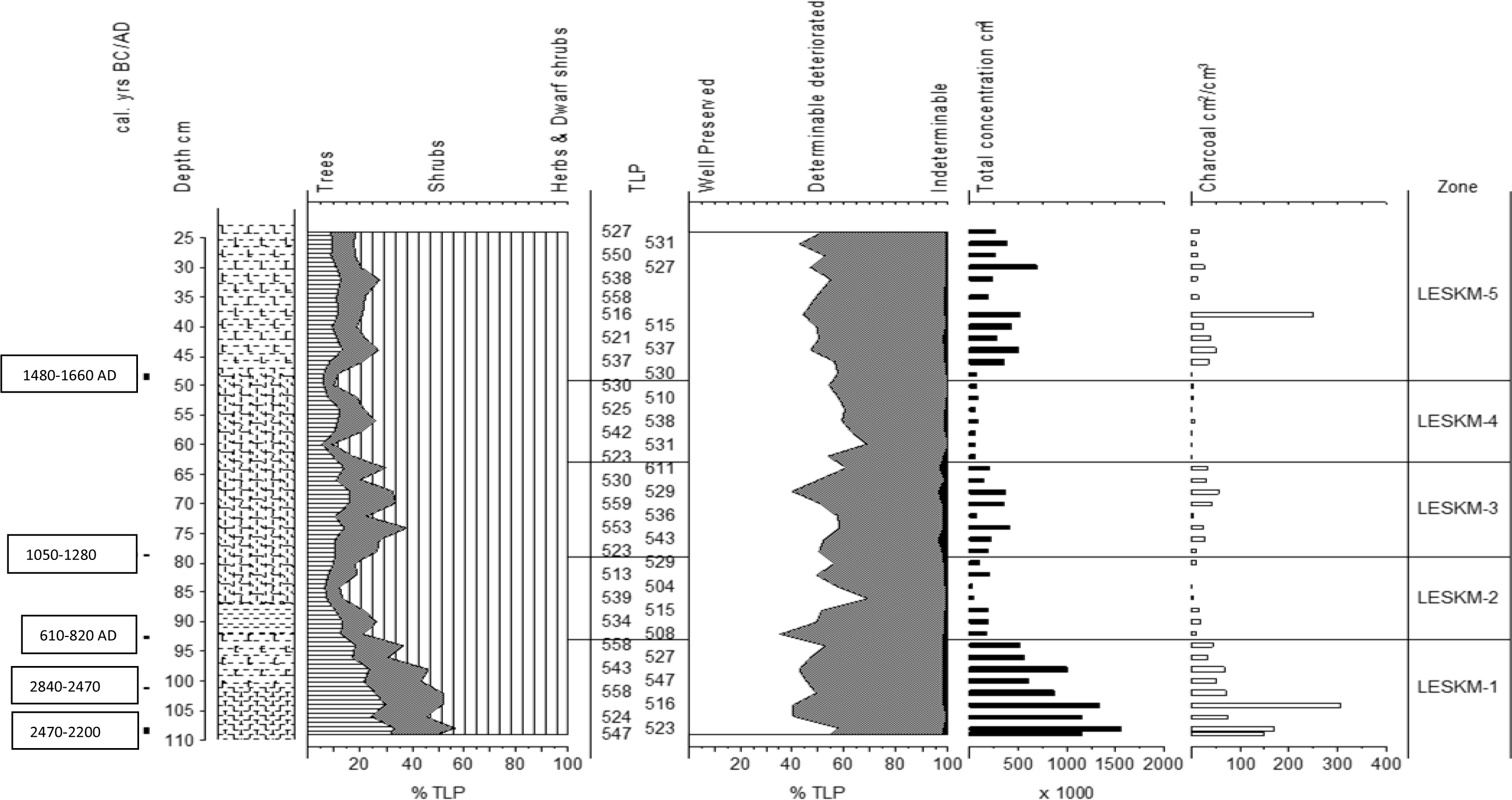
The Leskernick off-site pollen sequence: summary data, pollen concentrations and charcoal frequencies. For lithology see Table 1.

**Figure 5:**
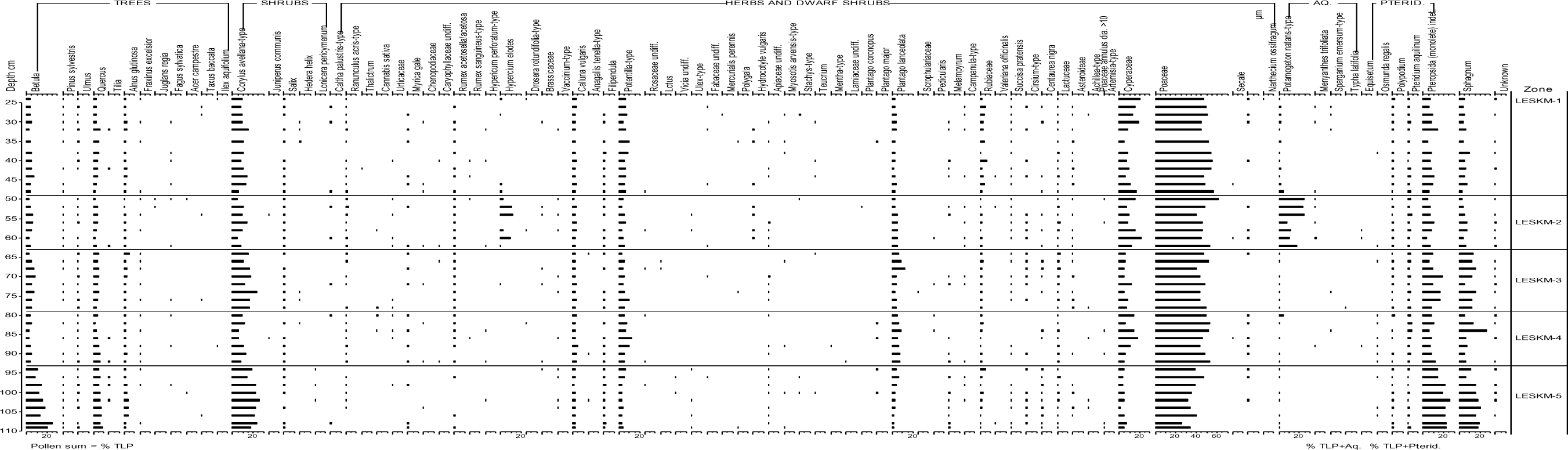
The Leskernick off-site pollen sequence. Percentage data for all taxa

#### Off-site pollen assemblage zones

##### LESKM-1 109-93 cm (Poaceae, Corylus avellana-type, Betula, Quercus zone)

Herbaceous pollen forms >45% TLP. Values for Poaceae, the main taxon, rise from *c*. 25 % TLP at the base to *c*. 45% TLP at the top of the zone. Of the trees and shrubs represented *Corylus avellana*-type (*c*. 20% TLP) and *Quercus* (*c*. 6% TLP) percentages are relatively stable, while those for *Betula* decline (from *c*. 20% TLP to *c*. 10% TLP). Pteropsida (monolete) indet. and *Sphagnum* values, initially *c*. 20% TLP+Group, also fall towards the top of the zone. Well preserved grains are generally in the minority (*c*. 40 % TLP) though the proportion of indeterminable grains is small (<2% TLP). Pollen concentrations for all groups decline from 1,560,000 cm^3^ to 520,000 cm^3^. Particularly high charcoal frequencies (>100 particles cm^2^/cm^3^) are recorded in some of the basal samples.

##### LESKM-2 93-79 cm (Poaceae, Cyperaceae, Potentilla-type zone)

Herbaceous pollen exceeds 70% TLP and rises to a maximum 88% TLP mid-zone. Poaceae remains the dominant taxon. Cyperaceae (maximum 18% TLP), *Potentilla*-type (maximum 11% TLP) and *Plantago lanceolata* (*c*. 7% TLP) are also well represented. Values for *Betula* and *Quercus* are generally <5% and *Corylus avellana*-type <15% TLP. Pteridophyte representation is also generally reduced compared to LESKM-1 as are *Sphagnum* values, though the latter rise to a peak of 28% TLP+*Sphagnum* towards the top of the zone. The proportion of well preserved grains rises to a maximum 70% TLP at 86 cm. Pollen concentrations and charcoal frequencies are low throughout the zone.

##### LESKM-3 79-63 cm (Poaceae, Corylus avellana-type zone)

This zone is characterised by higher tree and shrub pollen values. In particular *Corylus avellana*-type (maximum 24% TLP) and *Betula* (maximum 19% TLP) percentages increase. The representation of pteridophytes is also higher than in LESKM-2. Conversely values for the main herbaceous types are reduced (Poaceae to *c*. 45% TLP). The proportion of indeterminable grains rises to a maximum 3.5% TLP, though well preserved grains generally form >50 % TLP. Both pollen concentrations (maximum 415,000 cm^3^) and charcoal frequencies (maximum 56 particles cm^2^/cm^3^) are higher than in the preceding zone.

##### LESKM-4 63-49 cm (Poaceae, Cyperaceae, Potamogeton-natans, Hypericum elodes zone)

Poaceae, Cyperaceae and *Corylus avellana*-type pollen values return to percentages similar to those recorded in LESKM-2. High percentages are recorded for the bog herb *Hypericum elodes* (maximum 11% TLP) and the aquatic *Potamogeton natans*-type (maximum 25% TP). Pteropsida (monolete) indet. and *Sphagnum* values are consistently low. The proportion of well preserved grains reaches a maximum of 69% TLP at 60 cm. The pollen concentrations (*c*. 60,000 cm^3^) and charcoal frequencies (<5 particles cm^2^/cm^3^) are the lowest recorded from the monolith.

##### LESKM-5 49-24 cm (Poaceae, Cyperaceae, Corylus avellana-type, Potentilla-type zone)

The main taxa show little variation (Poaceae 45-50% TLP, *Corylus avellana*-type *c*. 10% TLP and *Potentilla*-type *c*. 5% TLP) with the exception of Cyperaceae, values for which exceed 10% TLP above 32 cm. Pteridophytes and *Sphagnum* percentages remain low. The proportion of well preserved grains declines from 58% TLP at the base to 52% TLP at the top of the zone. Pollen concentrations (*c*. 300,000 cm^3^) and charcoal frequencies are consistently higher than in the three preceding zones. Charcoal values peak (252 particles cm^2^/cm^3^) at 38 cm.

#### Chronology

A chronology for the off-site pollen investigations is provided by five AMS radiocarbon dates (Table 2). These suggest a hiatus in the lower part of sequence, which on the basis of the changes in biostratigraphy (including the pollen concentration data), charcoal frequencies and a change in lithology, is likely to be at the LESKM-1/2 boundary (at 92 cm). The increase in minerogenic material preceding this boundary may represent soil development but seems insufficient to represent the >2000 year gap and some erosion is therefore likely to have occurred.

**Table 2:** Radiocarbon dates obtained from the monolith.

| Depth from surface (cm) | 14C Age BP ( ±1 σ) | Calibration 2σ | d13C (‰) | Sample Code | Material dates |
| --- | --- | --- | --- | --- | --- |
| 48-49 | 300 ± 40 | 1480-1660 cal. yrs AD | -27.3 | Beta-133749 | Dark brown highly humified peat |
| 78.5-79 | 820 ± 50 | 1050-1280 cal. yrs AD | -25.9 | Beta-142320 | Brown/black humified Sphagnum/rootlet peat |
| 92.5-93 | 1330 ± 50 | 610-820 cal. yrs AD | -33.2 | Beta-142319 | Grey organic sand |
| 101-101.5 | 4050 ± 30 | 2840-2470 cal. yrs BC | -27.7 | Beta-144350 | Black organic mud |
| 108-109 | 3870 ± 40 | 2470-2200 cal. yrs BC | -27.7 | Beta-133748 | Black organic mud with gravel fragments |

The lack of an age gradient and the slight age inversion between Beta-133748 and Beta-144350 may be the result of AMS dating an organic deposit which, prior humification, contained large quantities of root material. On the basis of the dates available LESKM-1 probably represents the late Neolithic/early Bronze Age. The overlying sequence provides a high resolution record from the middle Saxon period to the recent past. Linear interpolation would show the sediment accumulation rate accelerating towards the present (see Figure 6). However, the occurrence of taxa indicative of particularly wet conditions (*Potamogeton natans*-type and *Hypericum elodes*), combined with low pollen concentrations and charcoal frequencies, suggests LESKM-4 accumulated rapidly and that interpolated ages should be treated with caution.

**Figure 6:**
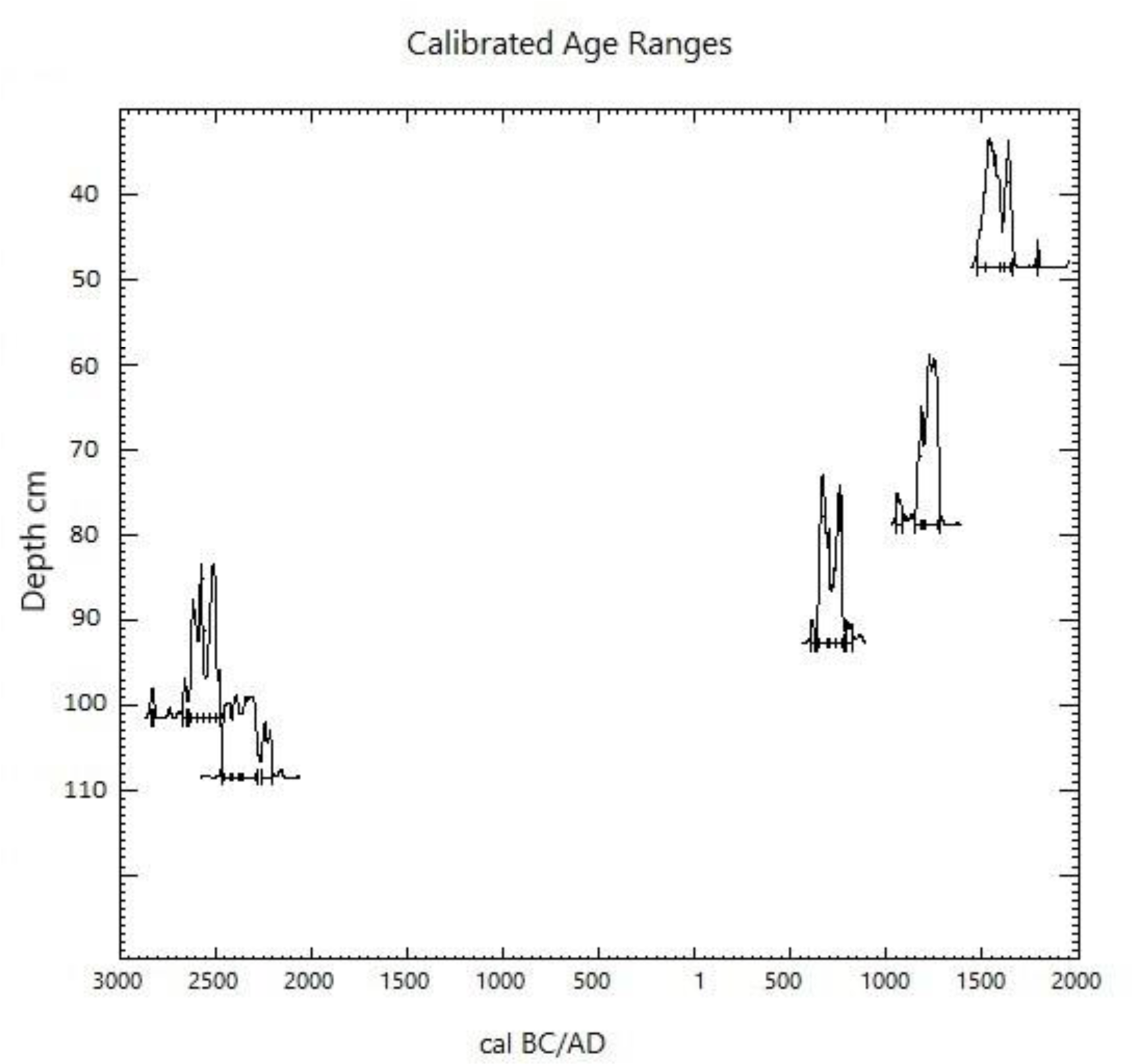
Age-depth plot for the Leskernick monolith, showing the probability distribution of the calibrated dates (from CALIB 8.1.0).

#### Interpretation of the off-site sequence

DCA analysis of the monolith samples was performed on all taxa that occurred at >4% TLP (Figure 7). The two taxa which achieve high values in LESKM-4, *Potamogeton natans*-type and *Hypericum elodes*, form a distinct outlying group in the DCA species plot (Figure 7a). As previously noted both are associated with wet places, though *Hypericum elodes* is tolerant of disturbance and is favoured by heavy grazing (Rodwell 1991). Axis 1 appears to distinguish wooded (e.g. *Quercus*, *Corylus avellana*-type, *Betula*, Pteropsida (monolete) indet. have high scores) from open habitats (e.g. Poaceae, Cyperaceae have low scores).

**Figure 7:**
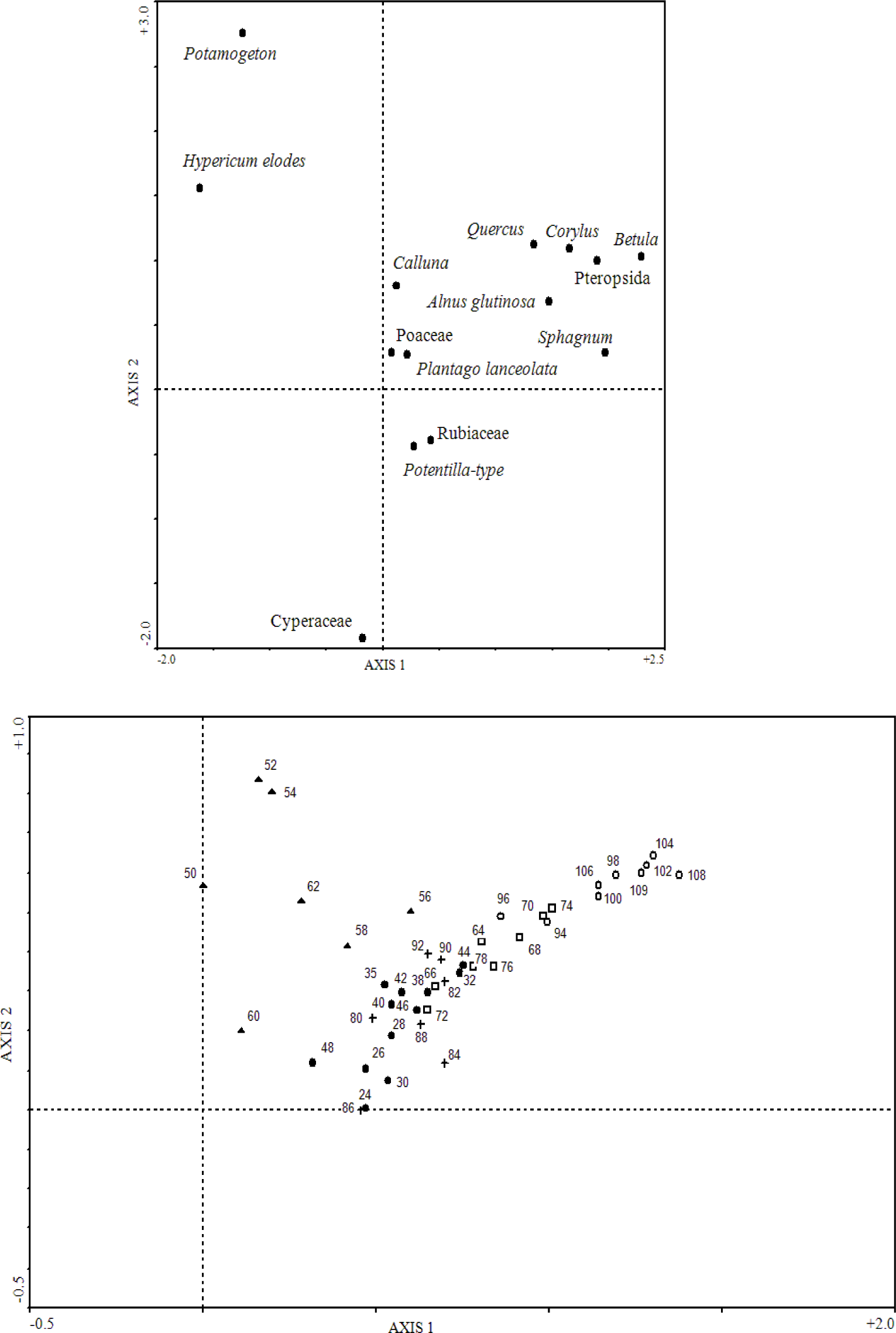
Plots of the DCA axis 1 and axis 2 scores from the off-site monolith. a species scores, b sample scores (LESKM-1 ○; LESKM-2 +; LESKM-3 □; LESKM-4 ▲; LESKM-5 ●)

The LESKM-1 assemblage is likely to have formed during the period the ritual complex at the base of the hill was constructed. It suggests a relatively open landscape, with some areas of *Corylus avellana*-type and *Betula* shrub/woodland. Given that Gearey *et al*. (2000a) report percentages of *Corylus avellana*-type of over 90% TLP at Rough Tor in the early Holocene, the much lower values in LESKM-1 assemblage suggest that the area around Leskernick had been subject to some prior anthropogenic disturbance. The changes in hydrology associated with this activity may well have been responsible for peat initiation at the site of the monolith (Moore 1975; Moore 1985). Trees/shrubs are most likely to have persisted on the heavily clittered slopes nearby. The local presence and open nature of the woodland remaining is indicated by the persistent occurrence of *Melampyrum*, which prefers glades and the pollen of which is poorly dispersed (Moore *et al*. 1986), and by the occurrence in LESKM-1 of a number of tall herbs (*Succisa pratensis*, *Centurea nigra*, Apiaceae) associated with woodland margins. *Melampyrum* is also associated with burning (Caseldine and Hatton 1993), which the high charcoal values confirm was occurring during the formation of the LESKM-1 assemblage. In the sequences of Gearey *et al*. (2000a) *Betula* is associated with taxa indicative of damp soils and the *Betula* pollen in LESKM-1 may have derived from woodland cover remaining in the wetter areas, though in contrast to Rough Tor and East Moor areas, *Alnus glutinosa* was not an important component of this community at Leskernick.

In addition to the evidence for burning, the LESKM-1 assemblage contains elements characteristic of grazing. *Plantago lanceolata*, *Potentilla*-type and Rubiaceae are all consistently present and increase towards the top of LESKM-1 suggesting an expansion in open habitats. Although predominately pastoral activity can be inferred, as with the Bronze Age sequences of Gearey *et al*. (2000b), sporadic cereal grains (Poaceae >10µ annulus diameter) occur. Cereal pollen is very poorly dispersed and their presence suggests small-scale cultivation was occurring close-by during the late Neolithic/early Bronze Age.

The lower axis 1 scores (Figure 7b) indicate woodland cover was reduced when peat formation recommenced (*c*. 700 cal. yrs AD). Indicators of human activity in the LESKM-2 assemblage include high pollen values for herbs indicative of grazing, particularly *Plantago lanceolata* and *Potentilla*-type. The latter is likely to have been derived from *Potentilla erecta* the abundance and flowering of which is greatly enhanced by grazing (Moore *et al*. 1986). Regular occurrence of cereal grains, at roughly the same time that Cerealia pollen is first recorded in the Stuffle pollen profile from Colliford (Walker in Austin *et al*. 1989), demonstrates that cultivation was occurring on Bodmin Moor. The early Medieval appears then to have been a period of comparatively intense human activity at Leskernick (in apparent contradiction to the suggestion of transhumance). However, comparison with LESKM-1, in terms of the extent of human activity may be misleading as the vegetation which produced the LESKM-2 assemblage will have been influenced by processes, both natural and anthropogenic, operating over the intervening years, some which are likely have had long-term effects on the environment. Soil deterioration and the lack of seed sources could, for example, have influenced the ability of trees/shrubs to regenerate irrespective of the level of grazing in the early Medieval period.

The LESKM-3 samples generally have higher axis 1 scores than those from LESKM-2 (Figure 7b) suggesting a late Medieval (from 1050-1280 cal. yrs AD) reduction in human activity in the Leskernick area. Both the local cessation of cultivation and woodland regeneration, the latter possibly over a larger area, are indicated. Reduced grazing pressure seems likely with tall herbs (including *Succisa pratensis*, *Centaurea nigra* and *Cirsium*-type) more consistently represented during LESKM-3. These findings appear to contrast with the previous work at Rough Tor and East Moor (Gearey *et al*. 1997, 2000a), and other sites in SW England (e.g. Austin *et al*. 1980; Geary *et al*. 1997). There is, however, evidence at Colliford of a contemporary decline in pastoralism and some woodland regeneration (Walker in Austin *et al*. 1989). The absence of cultivation in the immediate vicinity, may account for the lack of cereal pollen at Leskernick, with precise age of the nearby arable area shown by Herring *et al*. (2008) unknown. The apparent failure (within the constraints of the available chronology) of the Leskernick sequence to conform with the Medieval ‘high tide’, may then reflect both the local nature of the pollen record and the complexity of the changes occurring.

Increased human activity is indicated at Leskernick prior to 1480-1660 cal. yrs AD by a rise in *Plantago lanceolata*, the reappearance of cereal pollen and low *Corylus avellana*-type values at the top of LESKM-3 and during LESKM-4. However, as previously noted, LESKM-4 is characterised by high *Hypericum elodes*, *Potamogeton natans*-type and Cyperaceae pollen values. Increases in the first two taxa are consistent with the development of open pools on the valley floor with *Hypericum elodes* associated with areas where there is some water movement (Rodwell 1991). This increase in wetness could be considered to be only of local significance, a result of disturbance or representing a lateral change in the pattern of water seepage. However, peaks also occur in *Potamogeton* pollen in both the Stuffle (Walker in Austin *et al*. 1989) and Okehampton (Austin *et al*. 1980) sequences. Although poorly chronological constrained (dated to 1180-1410 cal. yrs AD, and between 1400-1640 cal. yrs AD and 1500-1890 cal. yrs AD, respectively), Austin *et al*. (1980) implicate climatic deterioration following the onset of the Little Ice Age. The high *Hypericum elodes* values are suggestive of disturbance and likely to represent the use and poaching of the expanded wet areas by grazing animals.

The drier conditions, indicated by lower aquatic pollen values during the formation of the post-medieval (1480-1660 cal. yrs AD onwards) LESKM-5 assemblage may reflect hydrological changes resulting from the onset of the tin industry. The lower *Plantago lanceolata* values suggest a decline in grazing pressure (from the late Medieval period). The consistently high values for *Potentilla*-type and Rubiaceae (likely to have been derived from *Galium saxatile*) appear contradictory. They could, however, be indicative of nearby spoil or, given that their pollen is poorly dispersed and the drier conditions, be derived from the bog surface. The consistency of the assemblage is surprising with the vegetation stability implied suggesting the assemblage formed before the major changes in the valley bottom brought about by tin streaming, which presumably resulted in the development of the modern soil (the uppermost 25 cm was not sampled).

### On-site investigations

#### Introduction and taphonomic considerations

The on-site investigations were undertaken from a variety of contexts to address questions that arose during the archaeological excavations. While the taphonomy of these assemblages is in some cases straight forward (e.g. the buried peats) in most cases, the buried land surfaces/turf lines and their underlying soils, it is likely to be more complex. It is often assumed that the pollen extracted from such material will provide information on the composition of the local vegetation immediately prior to burial. Active (mull) soils appear to rapidly incorporate pollen grains, largely through the mixing action of earthworms (Andersen 1979b; Aaby 1983; Davidson *et al*. 1999). Decay is, however, also rapid and past pollen spectra should quickly become diluted by continued mixing with new pollen input, until the soil is sealed (Tipping 1994).

There are a number of potential problems. Firstly the presence of residual older pollen which may have survived in the soil for some time prior to burial. Models of pollen survival derived from mull soils are likely to be inappropriate in the case of the acidic and wet soils of Bodmin Moor, where residual pollen may make up a high proportion of the total rather than being swamped by fresh pollen incorporated immediately prior to burial.

A second issue is the downward movement of more recently released pollen, if a buried soil is overlain by porous deposits. However, an investigation into this problem by Davidson *et al*. (1999) found no evidence of significant down-washing, despite examining a number of soil types subject to variety of hydrological regimes.

In addition, pollen assemblages from mor humus and podzol profiles have been used, not without controversy (e.g. Godwin 1958), to infer vegetation changes over time (see Dimbleby 1985). Stratification is thought to occur as pH and biological activity gradually decline, leaving pollen assemblages in chronological order. Work by Tipping *et al*. (1999), however, indicates that in such soils pollen is incorporated and mixed into the near surface organic horizons over long time periods and that such layers are unlikely to contain stratified assemblages. There was little evidence for the movement of pollen into the underlying mineral horizon. The latter may then preserve pollen from an earlier more active phase, though these authors caution that the temporal resolution of podzol profiles is likely to be very poor.

The state of preservation of pollen grains can be used provide evidence of pollen ‘contamination’ from adjacent sediment (e.g. the presence of large quantities of ‘fresh’ or severely deteriorated pollen) and the ‘reliability’ of the pollen spectra. The on-site assemblages at Leskernick are generally poorly preserved and may therefore have undergone considerable post-depositional alteration, being biased towards taxa which are either, robust and preferentially preserved, or are readily identifiable in a deteriorated state. Vegetation reconstructions based on such data will inevitably be flawed. To identify samples that may have been subject to such alteration, the recommendations of Bunting and Tipping (2000) have been followed. Bunting and Tipping (2000) identified 9 criteria against which samples can be evaluated, to determine whether they may have been subject to post-depositional bias. They caution against adopting any single criteria, particularly high percentages of ‘robust’ types as several of these are common in agricultural environments. In general, using these criteria, the on-site Bodmin assemblages show few signs of post-depositional alteration (Table 3). Four samples (Table 3) exceed the thresholds in more than two categories and can therefore, following Bunting and Tipping (2000), be considered unreliable. These samples (BSA98-1 38 cm, LSW1-44 27, 30, 34 cm) are shaded on the relevant pollen diagrams.

**Table 3:** On-site pollen samples which exceed the thresholds in the property categories of Bunting and Tipping (2000). The pollen types included under the resistant category are *Tilia*, Caryophyllaceae, Lactuceae, *Artemisia*-type and Brassicaceae. Only Lactuceae was recorded at values >6% TLP.

| Total land pollen sum <300 | Total pollen concentration <3000 cm<sup>3</sup> | Number of taxa in main sum <10 | Percentage degraded >35% | Percentage indeterminate >30% | Percentage resistant >6% (Lactuceae) | Pteropsida (monolete) indet. >40% | Spore: pollen concentration ratio >0.66 | Spore: pollen taxa ratio >0.66 |
| --- | --- | --- | --- | --- | --- | --- | --- | --- |
| All samples exceed | All samples exceed | All samples exceed | LNC-27 29, 41, 56 cm<br>BSE99 55 cm<br>BSD99-4 22, 24 cm<br>BEB99 10 cm | No samples exceed | LSR-14A 31 cm<br>LSR-14B 27 cm<br>LSR-27 9, 14 cm<br>BSA98-1 33, 38 cm<br>BSB98-2 9, 16 cm<br>BEB99 17, 20 cm | LSW23-7 1, 2, 3, 4, 5 cm<br>LSW1-44 27, 30.5, 34 cm | LSW23-7 0 cm<br>LSW1-44 27, 30.5, 34 cm<br>LSW1-33 34, 37 cm<br>L<br>BSA98-1 35, 38 cm<br>LSS39-11 15 cm<br>BSB99-B | No samples exceed |

The results from the on-site contexts which relate to the ritual features (a Stone Circle and Stone Row) are presented first. These are followed by the samples from houses/huts (usage here follows that of the excavators) in the Western and Southern Settlements and finally those collected from beneath boundary sections and entrances (see Figure 1). Radiocarbon dates obtained from the features investigated palynologically are shown in Table 4. A full list of the dates obtained during the archaeological excavations can be found in Bender *et al*. (2007). These indicate human activity in the vicinity of the settlement from *c*. 1600 to 600 cal. BC.

**Table 4.**
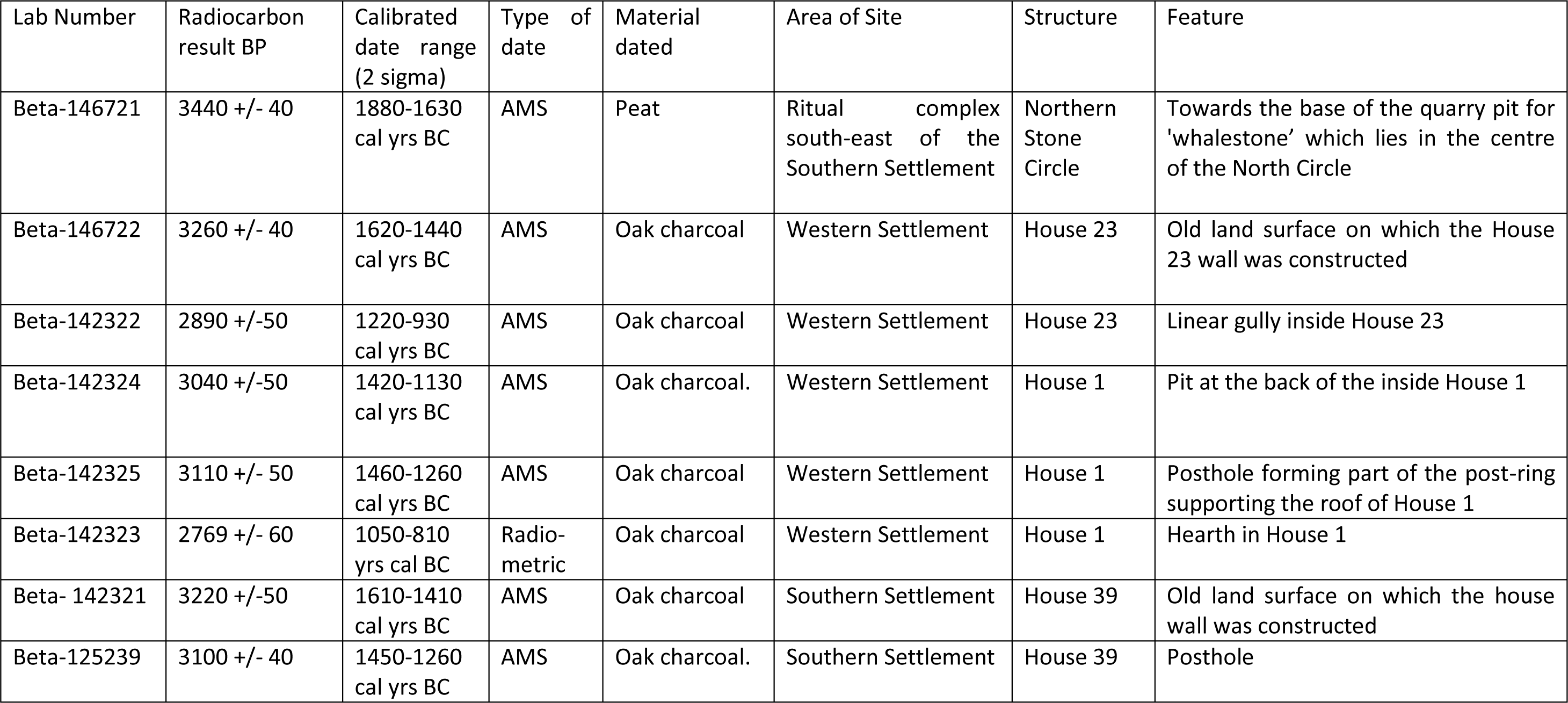
Leskernick radiocarbon dates related to structures from which pollen samples were collected (from Bender *et al*. 2007) recalibrated using CALIB 8.1.0.

| Lab Number | Radiocarbon result BP | Calibrated date range (2 sigma) | Type of date | Material dated | Area of Site | Structure | Feature |
| --- | --- | --- | --- | --- | --- | --- | --- |
| Beta-146721 | 3440 +/- 40 | 1880-1630 cal yrs BC | AMS | Peat | Ritual complex south-east of the Southern Settlement | Northern Stone Circle | Towards the base of the quarry pit for 'whalestone' which lies in the centre of the North Circle |
| Beta-146722 | 3260 +/- 40 | 1620-1440 cal yrs BC | AMS | Oak charcoal | Western Settlement | House 23 | Old land surface on which the House 23 wall was constructed |
| Beta-142322 | 2890 +/-50 | 1220-930 cal yrs BC | AMS | Oak charcoal | Western Settlement | House 23 | Linear gully inside House 23 |
| Beta-142324 | 3040 +/-50 | 1420-1130 cal yrs BC | AMS | Oak charcoal. | Western Settlement | House 1 | Pit at the back of the inside House 1 |
| Beta-142325 | 3110 +/- 50 | 1460-1260 cal yrs BC | AMS | Oak charcoal | Western Settlement | House 1 | Posthole forming part of the post-ring supporting the roof of House 1 |
| Beta-142323 | 2769 +/- 60 | 1050-810 yrs cal BC | Radio-metric | Oak charcoal | Western Settlement | House 1 | Hearth in House 1 |
| Beta- 142321 | 3220 +/-50 | 1610-1410 cal yrs BC | AMS | Oak charcoal | Southern Settlement | House 39 | Old land surface on which the house wall was constructed |
| Beta-125239 | 3100 +/- 40 | 1450-1260 cal yrs BC | AMS | Oak charcoal. | Southern Settlement | House 39 | Posthole |

#### The Northern Stone Circle

The Northern Stone Circle (LNC) lies at the base of the hill with the Stone Row (LSR) nearby (Figure 1). At LNC pollen investigations were undertaken from the organic fill of a stone hole (described as peaty silt with stones, context 15) and an overlying organic deposit (‘peat’). The stone (referred to as a ‘whalestone’) had been dug out from a central location and was lying partially across the hole (Bender *et al*. 2007). An Early Bronze Age radiocarbon date (1880-1630 cal. yrs BC) was obtained from the peaty silt indicating that stone was extracted in prehistory. The pollen samples were collected partially to determine whether the hole had been back-filled when the stone was extracted or whether the hole was left open and gradually became infilled with the peaty silt and stones. Five pollen assemblage zones (prefixed LNC) have been defined (Figure 8).

**Figure 8:**
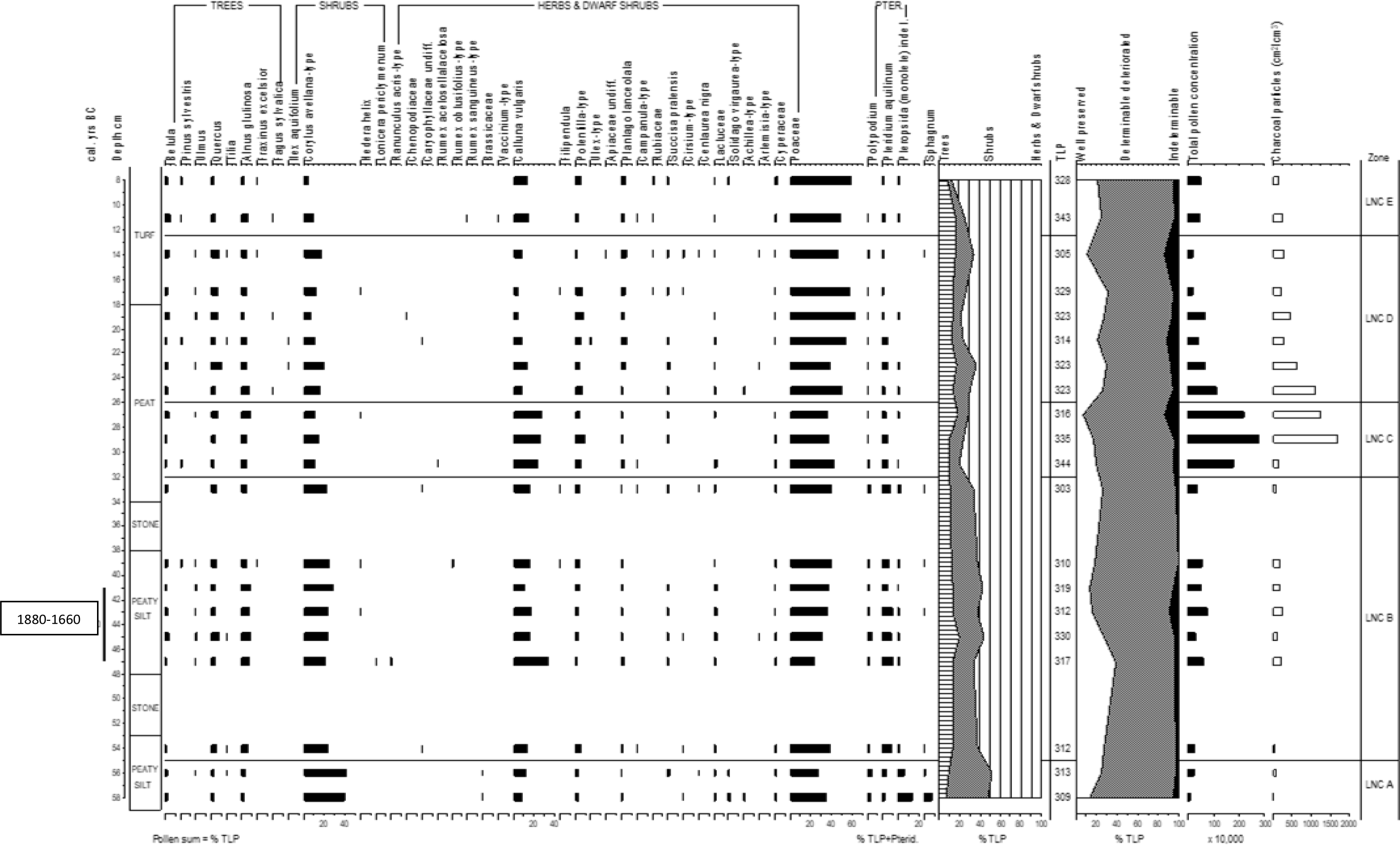
The Northern Stone Circle (LNC). Percentage pollen data, summary data, pollen concentrations and charcoal frequencies.

LNC A 58-55 cm (Corylus avellana-type, Poaceae, Calluna vulgaris, Pteropsida (monolete) indet. zone)

*Corylus avellana*-type values of *c*. 40% TLP are recorded, though pollen from other trees and shrubs is scarce. Poaceae (maximum 35% TLP) and *Calluna vulgaris* (maximum 12% TLP) are the other main taxa. Pteropsida (monolete) indet. spores decline from 13% to 6% TLP+Pterid. Over 70% TLP of the assemblage is poorly preserved. Pollen concentrations are <260,000 cm^3^ and charcoal frequencies <80 cm^2^/cm^3^.

##### LNC B 55-32 cm (Poaceae, Corylus avellana-type, Calluna vulgaris, Alnus glutinosa, Pteridium aquilinum zone)

*Corylus avellana*-type values are *c*. 25% TLP, though the representation of both *Alnus glutinosa* (maximum 8% TLP) and *Quercus* (maximum 7% TLP) is higher than in the preceding zone. *Calluna vulgaris* values are also higher, peaking (34% TLP) at 47 cm. *Pteridium aquilinum* values of *c*. 9% TLP+Pterid. are recorded. The percentage of well-preserved grains declines from 38% to 13% TLP between 47 and 41 cm. Pollen concentrations of a maximum 777,000 cm^3^ and charcoal frequencies of a maximum 250 cm^2^/cm^3^ are recorded.

##### LNC C 32-26 cm (Poaceae, Calluna vulgaris, Corylus avellana-type, Potentilla-type zone)

This zone is characterised by relatively high *Calluna vulgaris* (*c*. 26% TLP) and low *Corylus avellana*-type (*c*. 11% TLP) values. *Potentilla*-type attains a maximum 10% TLP. Pollen preservation declines to a minimum of 6% TLP well preserved at 27 cm. Both pollen concentrations and charcoal frequencies peak during this zone attaining maximum values (2,198,000 cm^3^ and 1,700 cm^2^/cm^3^, respectively) at 29 cm.

##### LNC D 26-12 cm (Poaceae, Corylus avellana-type, Calluna vulgaris, Quercus zone)

Poaceae pollen dominates this zone rising to a maximum 63% TLP at 19 cm. *Calluna vulgaris* values (<14% TLP) are much reduced from the preceding zones, while *Corylus avellana*-type values are similar and those for *Quercus* increase (maximum 10% TLP). Well preservation grains make up *c*. 30% TLP. Pollen concentrations and charcoal frequencies decline upwards to *c*. 200,000 cm^3^ and *c*. 250 cm^2^/cm^3^, respectively.

##### LNC E 12-8 cm (Poaceae, Calluna vulgaris zone)

Herbaceous pollen dominates this zone (maximum 88% TLP). Poaceae is the main taxon though *Calluna vulgaris* values of *c*. 14% TLP are recorded. *Corylus avellana*-type frequencies decline to *c*. 5% TLP. Well preserved grains make up *c*. 20% TLP, pollen concentrations of *c*. 500,000 cm^3^ and charcoal frequencies of *c*. 200 cm^2^/cm^3^ are recorded.

The occurrence of two distinct pollen assemblages and the consistent pollen stratigraphic changes (e.g. the rise in Poaceae in LNC B) within the peaty silt fill of the stone hole suggests that the hole was initially left open to accumulate sediment after the stone was extracted.

The LNC A assemblage contains the highest *Corylus avellana*-type frequencies recorded from Leskernick indicating the occurrence of hazel woodland/shrub on the surrounding hillsides when the stone was extracted. In contrast *Quercus* and *Alnus glutinosa* values are low. The LNC B is difficult to interpret. The decline in *Corylus avellana*-type and increases in *Pteridium aquilinum* and *Calluna vulgaris* suggest some woodland clearance, though this assemblage seems not to be indicative of a phase of intensive land use, particularly given that increases occur in *Quercus* and *Alnus glutinosa* pollen. Interestingly, the climbers *Hedera helix* and *Lonicera periclymenum* are both recorded in LNC-B. The pollen of these taxa is poorly dispersed and there occurrence supports the view that following the extraction of the stone, at least in the vicinity of the stone circle, human activity/grazing pressure was lax. The rises Poaceae and *Potentilla*-type towards the top of LNC B are indicative of increased grazing pressure.

The sequence above the infill is much thicker than that recorded in a control pit (LSR-CP) nearby and contains three distinct assemblages. LNC C contains high pollen concentrations and charcoal frequencies, which combined with the poor preservation, suggest that it may have accumulated over a long period of time. The combination of the high *Calluna vulgaris* and charcoal frequencies indicates periodic burning. Subsequently the grazing regime may have intensified with both Poaceae and *Plantago lanceolata* values increasing in LNC D. The upper assemblage with high Poaceae and *Calluna vulgaris* is comparable with the assemblage from the turf in the control pit (LSR-CP).

### Stone Row

#### 1) Samples from a control pit

Samples were collected from a control pit, dug adjacent to the Stone Row (LSR), to determine the pollen content of the modern turf and underlying soil, for comparison with sequences which may have been sealed as a consequence of past human activity. The samples (0 cm is the modern surface) from 5, 9 and 11 cm are from the modern turf and those from 14 and 15 cm are derived from a leached (Ea) horizon. One LPAZ (LSR-CP) has been defined (Figure 9).

**Figure 9:**
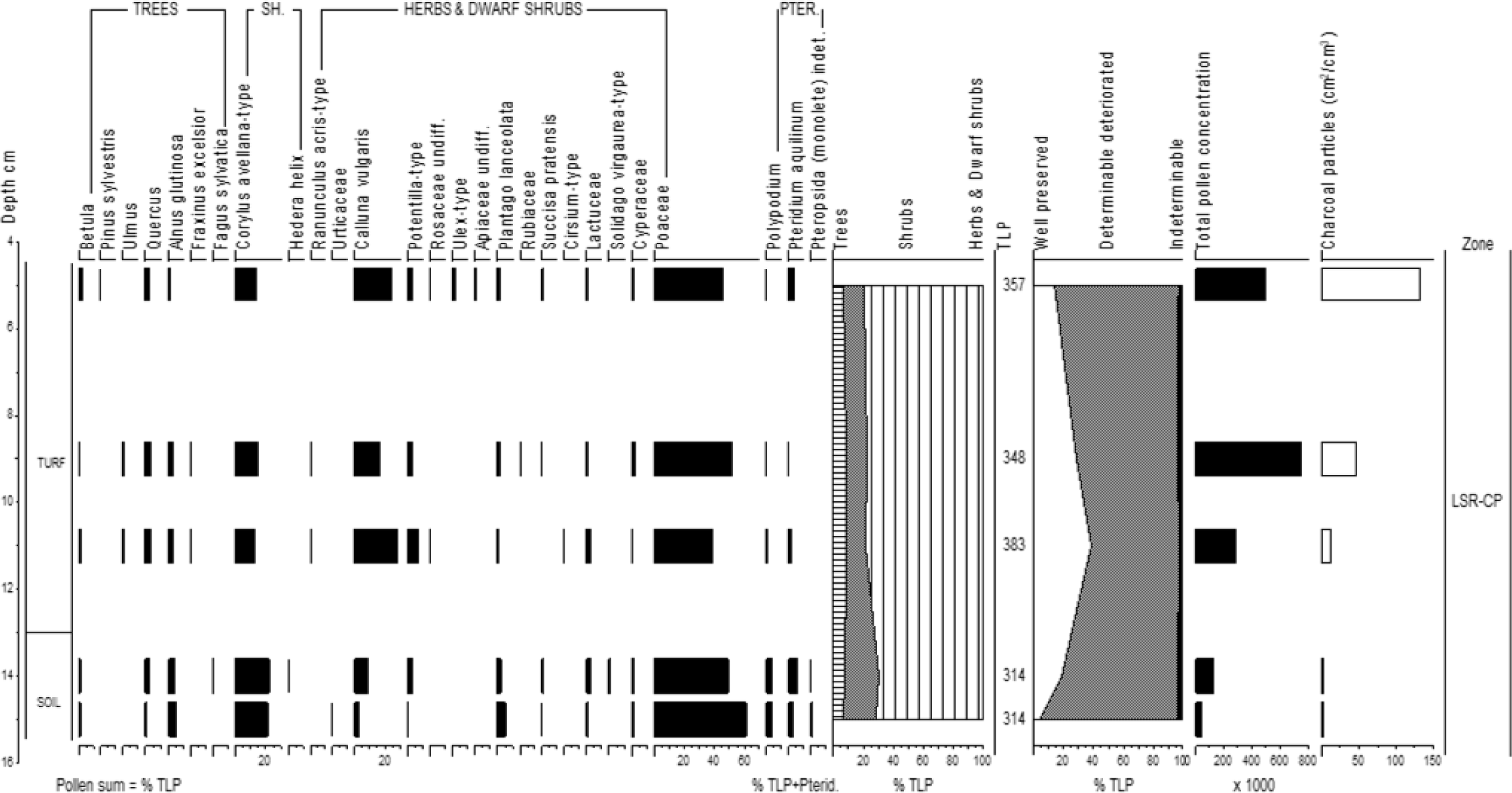
Samples from a control pit adjacent to the Stone Row (LSR-CP). Percentage pollen data, summary data, pollen concentrations and charcoal frequencies.

##### LSR-CP 15-5 cm (Poaceae, *Calluna vulgaris*, *Corylus avellana*-type zone)

Poaceae pollen (39-61% TLP) is dominant throughout. The values for *Calluna vulgaris* are notably higher in the turf than the leached horizon, while the reverse is the case for *Corylus avellana*-type and *Plantago lanceolata*. The proportion of grains well preserved peaks (39% TLP) at 11 cm. The pollen concentrations (maximum 744,000 cm^3^) rise above 11 cm and the charcoal frequencies increase progressively towards the top.

Values for *Corylus avellana*-type in the turf seem high given the nature of the contemporary vegetation close to the Stone Row. This pollen may be residual, though *Corylus avellana*-type pollen is well dispersed and values are similar to those in the upper (LESKM-5) assemblage in the off-site diagram. The pollen assemblages from the turf and soil are distinct with the high *Calluna vulgaris* values in the turf reflecting its local abundance. The underlying leached horizon may therefore contain a greater proportional of residual pollen from an earlier period/s in which heathland was less prominent.

#### 2) Samples from fill beneath a prostrate stone (from the western terminal)

Two series of samples (A and B) were collected from beneath a stone (context 14) lying flat on the ground. Samples from 21 cm (A and B) are from the turf (context 1) beneath the stone (not completely sealed). Samples from 27 cm (A and B) are from a 1^st^ fill (context 5) of a cut (probably the stone socket). Samples from A at 31 cm and B at 28 and 33 cm are from a 2^nd^ fill (context 10) of this cut. The basal samples (A 36 cm and B 36 cm) are from the soil beneath the cut. The depths are relative to the upper surface of the stone.

#### Series A) One LPAZ has been defined (Figure 10)

##### LSR14A 36-21 cm (Poaceae, Calluna vulgaris, Corylus avellana-type zone)

Tree and shrub pollen, mostly notably *Corylus avellana*-type, declines upwards (from 28-12% TLP). Values for Poaceae increase (maximum 62% TLP) and *Calluna vulgaris* (maximum 19% TLP) decline, except for the uppermost sample where these trends are reversed. Pollen preservation improves from *c*. 25% TLP to 56% TLP well preserved in the uppermost sample. Pollen concentrations are consistently >1,000,000 cm^3^ except in the lower-most sample and charcoal frequencies are generally low, though increase in the top sample (to a maximum of 62 cm^2^/cm^3^).

**Figure 10:**
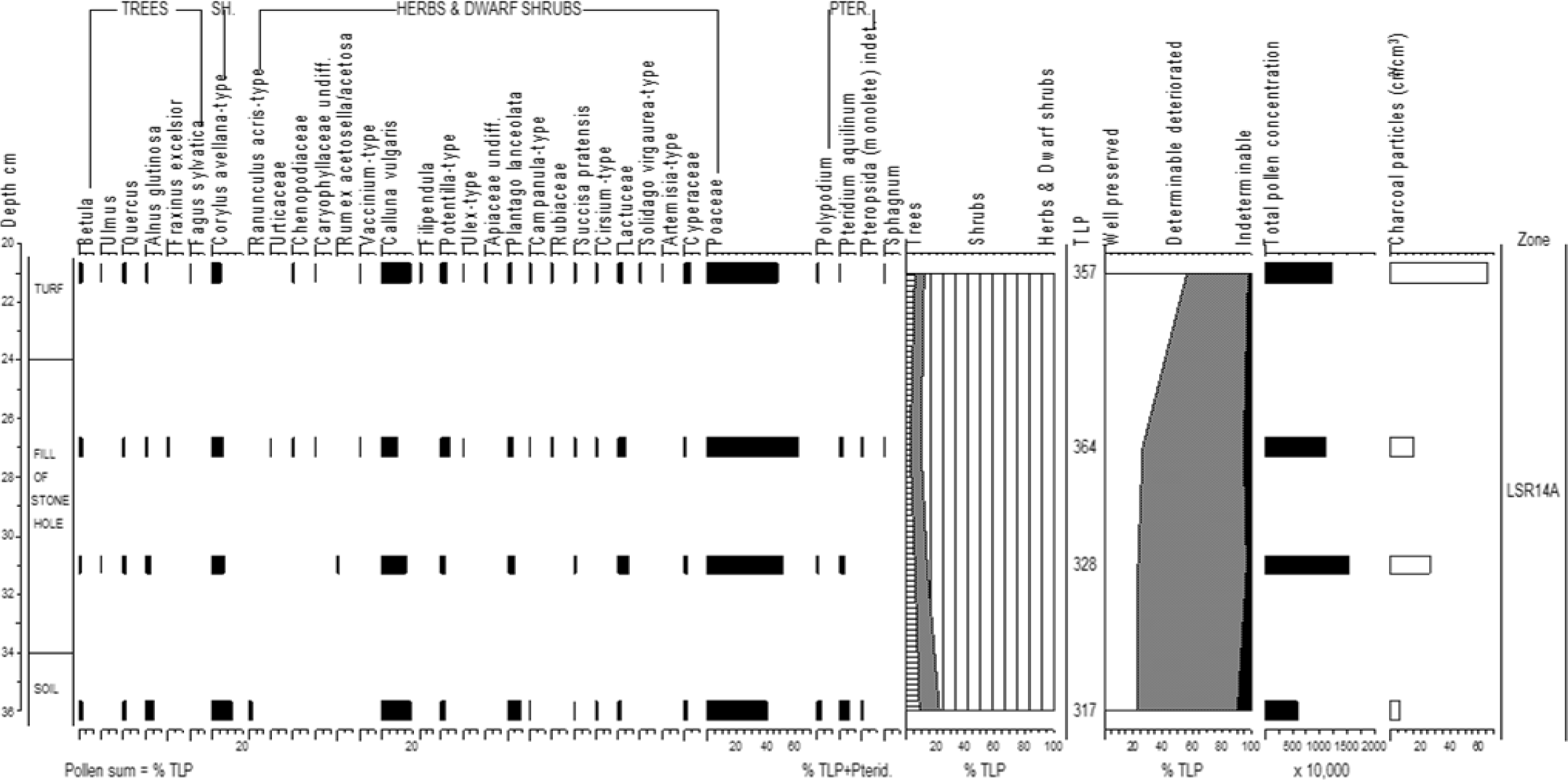
The first series of samples (series A) from fill beneath a prostrate stone associated with the Stone Row (LSR-14A). Percentage pollen data, summary data, pollen concentrations and charcoal frequencies.

#### Series B) One LPAZ has been defined (Figure 11)

##### LSR14B 36-21 cm (Poaceae, Calluna vulgaris, Corylus avellana-type zone)

An assemblage dominated by herbaceous pollen (>80% TLP). Poaceae percentages are higher above 33 cm (rising to *c*. 55% TLP) while values for *Calluna vulgaris* are lower. *Corylus avellana*-type values decline more steadily upwards. Well preserved grains form *c*. 15% TLP. Pollen concentrations are generally high (maximum 2,260,000 cm^3^ at the top) and follow trends in the charcoal frequencies, with the highest values recorded at 33, 28 and 21 cm.

**Figure 11:**
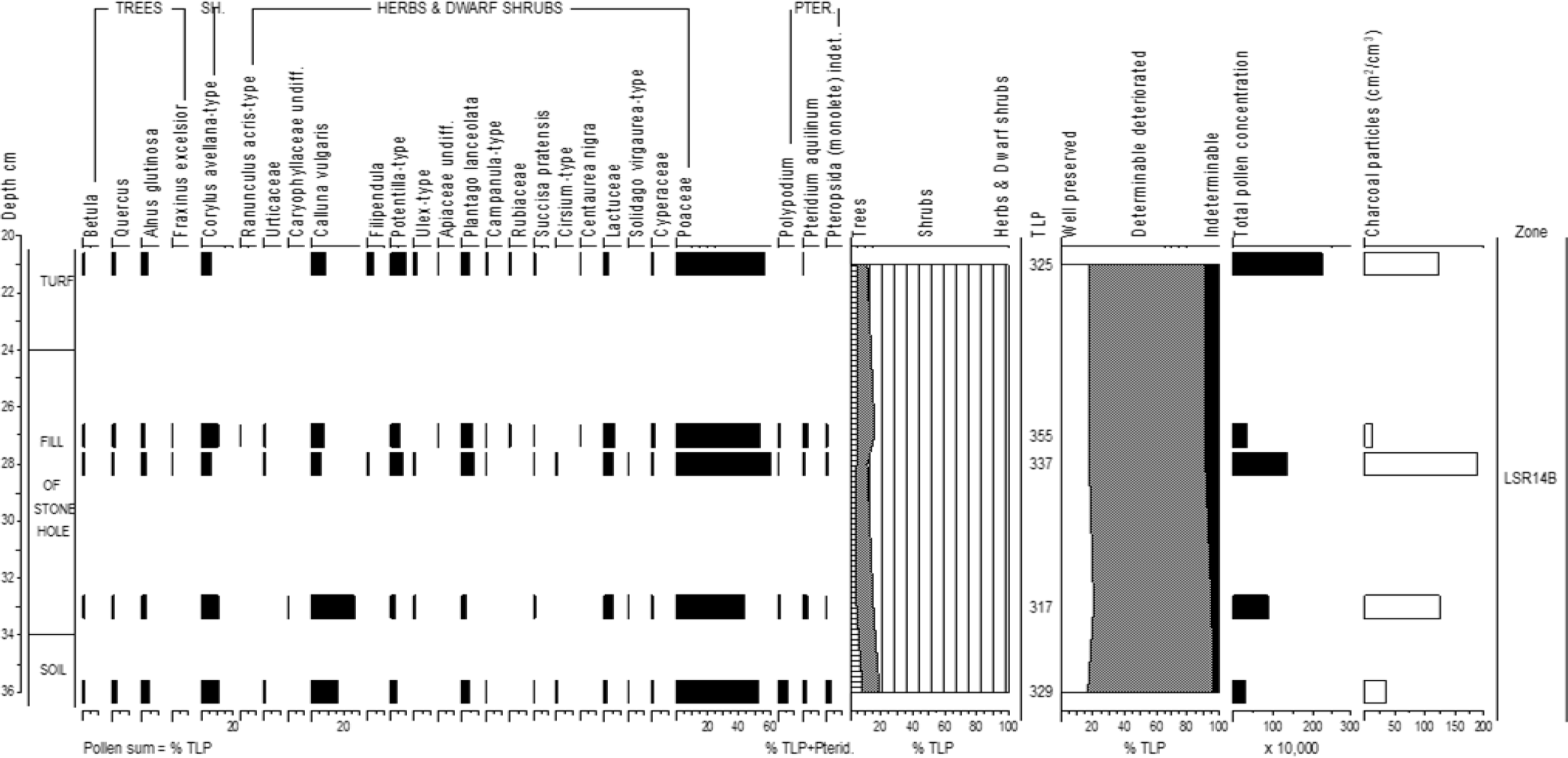
The second series of samples (B) from fill beneath a prostrate stone associated with the Stone Row (LSR-14B). Percentage pollen data, summary data, pollen concentrations and charcoal frequencies.

The pollen sequences from the two series are similar to each other and the different contexts do not contain particularly distinctive assemblages, though as would be expected pollen concentrations are much lower in the underlying soil. The similarity between the pollen content of the turf and fills suggests that much of the pollen in the fill sequences may have become incorporated after sediment deposition. Given that these assemblages are similar to those recorded from the turf in the control pit, which should be accumulating ‘fresh’ pollen, this may have occurred after the stone became prostrate, or indicate that this ‘dismemberment’ occurred relatively recently (see Bender *et al*. 2007).

#### 3) Samples beneath a spoil heap (Figure 12)

The largest stone in the row (the Great Terminal Monolith) was, at some point, dug out and pushed over. The spoil heaps from this activity seal a turf line and soil. Samples through the sequence were collected to determine whether the stone was extracted in the recent past. The sample (0 cm is the modern surface) from 5 cm is from an orange sand (the spoil, context 26).

**Figure 12:**
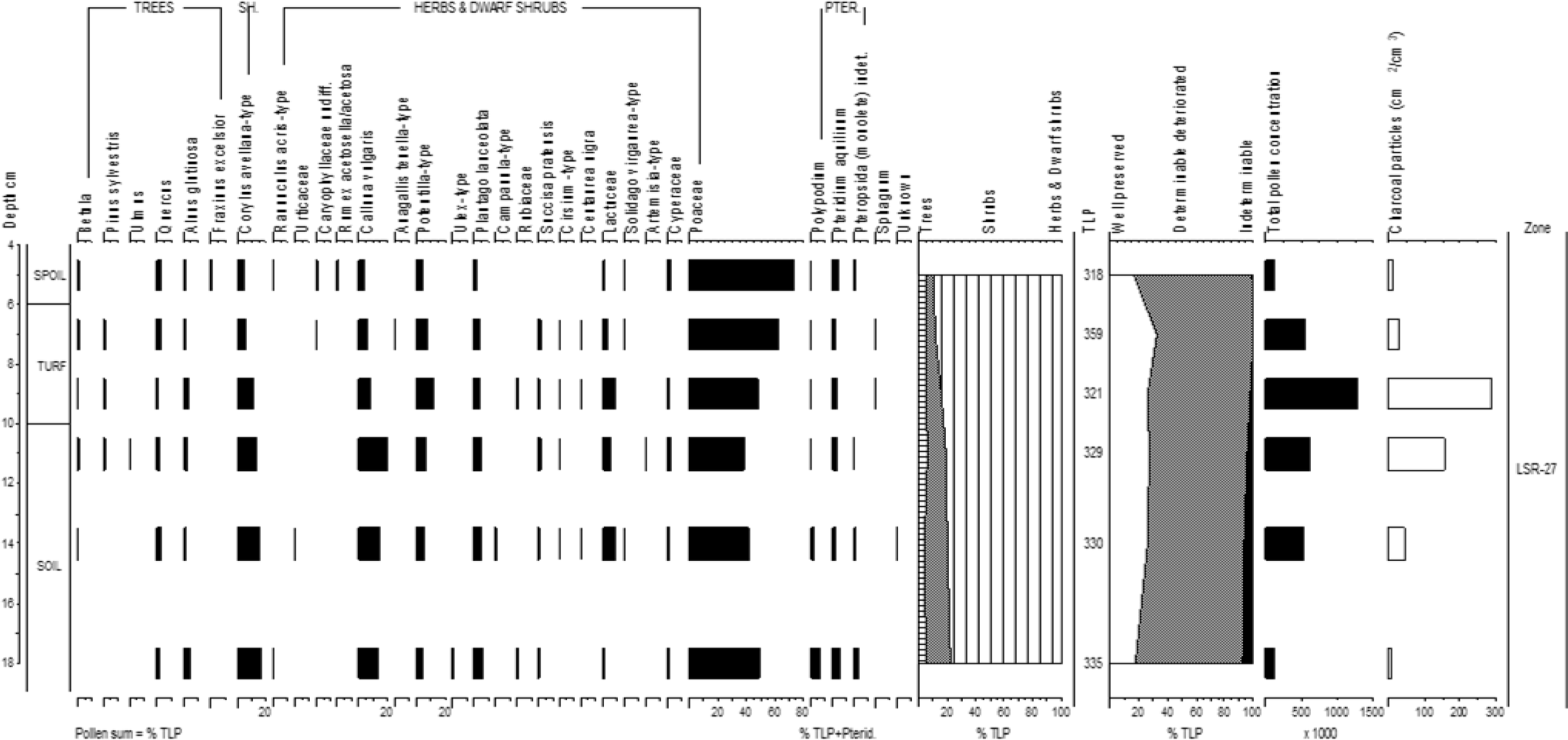
Samples beneath a spoil heap associated with the Stone Row (LSR-27). Percentage pollen data, summary data, pollen concentrations and charcoal frequencies.

### LSR-27 18-5 cm (Poaceae, Corylus avellana-type, Calluna vulgaris, Potentilla-type, Lactuceae zone)

Herbaceous pollen dominates these samples with Poaceae frequencies rising to a maximum 73% TLP at the top. Conversely *Corylus avellana*-type and *Calluna vulgaris* values decline upwards. *Potentilla*-type percentages are highest mid-zone where the pollen concentrations and charcoal frequencies peak (1,276,000 cm^3^ and 290 cm^2^/cm^3^, respectively). Well preserved grains form *c*. 25% TLP.

The high pollen concentrations and charcoal frequencies from 9 cm are consistent with the identification of a former turf line. The upper part of this assemblage appears distinct (more Poaceae and less *Calluna vulgaris*) from the control pit sequence (LSR-CP). The turf appears therefore have accumulated pollen and the spoil been excavated during a period in which grassland taxa were more abundant and heathland species less so, compared to today. Alternatively, the differences between the assemblages may indicate vegetation changes at a very local scale. As with the other assemblages from this area (LNC and LSR) *Corylus avellana*-type values are relatively high, particularly within the soil indicating they are probably in part residual.

### Western Settlement (LSW) House 23

‘House’ 23 is described as a small structure, less than 4m in diameter, possibly an open - air shrine, potentially in use over several hundred years (Bender *et al*. 2007).

#### 1) Samples from beneath stone 8 of the house wall

A Middle Bronze Age date of 1620-1440 cal. yrs BC (Table 4) was obtained (on *Quercus* charcoal) from the old land surface on which House 23 was constructed. These samples were taken from the land surface (0-2cm) and underlying soil (2-7 cm) beneath the house wall to investigate the environment prior to the construction of the house. The land surface showed signs of disturbance and may have been affected by small mammal activity. One LPAZ (LSW23-8) has been defined (Figure 13).

**Figure 13:**
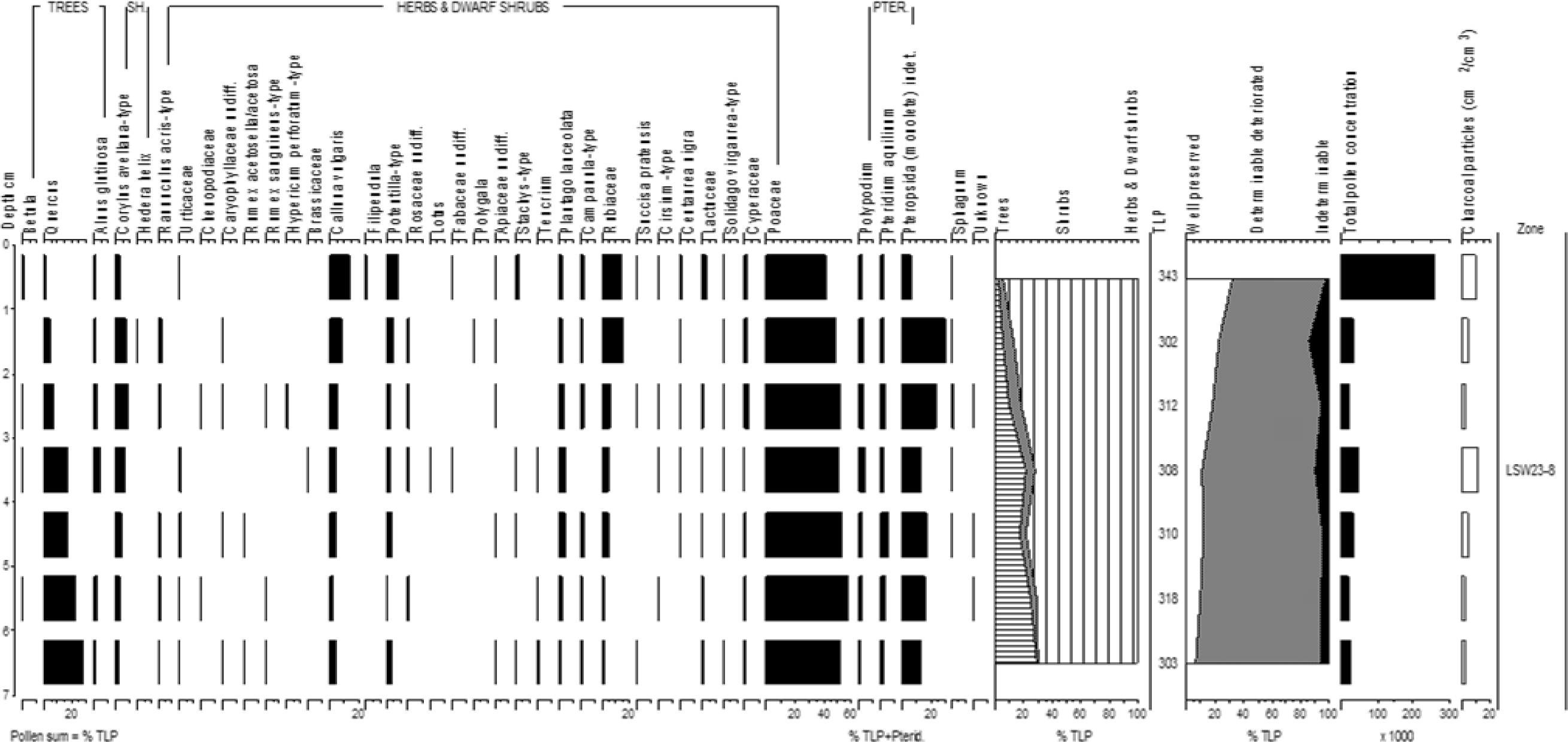
Samples from beneath stone 8 of the wall of House 23 West Settlement (LSW23-8). Percentage pollen data, summary data, pollen concentrations and charcoal frequencies.

##### LSW23-8 7-0 cm (Poaceae, Quercus, Calluna vulgaris, Rubiaceae zone)

High Poaceae values (>40% TLP) occur throughout. *Quercus* percentages decline steadily from the base (27% TLP) to the top (1% TLP), while those for *Calluna vulgaris*, Rubiaceae and *Potentilla*-type rise. *Corylus avellana*-type and *Plantago lanceolata* values peak mid-zone. Pteropsida (monolete) indet. spores exceed >12% TLP+Pterid. except in the uppermost sample. The proportion of well preserved pollen rises from 7% TLP at the base to 33% TLP at top of the assemblage. Pollen concentrations rise in the top sample from <45,000 grains cm^3^ to 258,000 cm^3^. Charcoal values are < 11 cm^2^/cm^3^ throughout.

The high proportion of well-preserved grains in the uppermost sample, combined with the high pollen concentrations and the indications of the disturbance of the land surface, suggest pollen could have become recently incorporated into the buried land surface. However, it seems more likely, given the gradational nature of the changes in pollen stratigraphy, that biological activity progressively declined prior to/or as a result of, the soil being sealed by the wall. If so then largely open conditions existed in the vicinity of the house immediately prior to its construction. The high basal *Quercus* pollen values may be residual from an earlier period, though the survival of the latter in the landscape is consistent with the presence of the Bronze Age *Quercus* charcoal.

#### 2) Samples from beneath stone 34 of the house wall

Investigations were also undertaken from beneath stone 34 to determine conditions prior to the construction of House 23. Samples (0-10 cm) were collected from the land surface and underlying soil (contexts 2 and 4). Both contexts may have disturbed by small mammal activity. One LPAZ (LSW23-34) has been defined (Figure 14).

**Figure 14:**
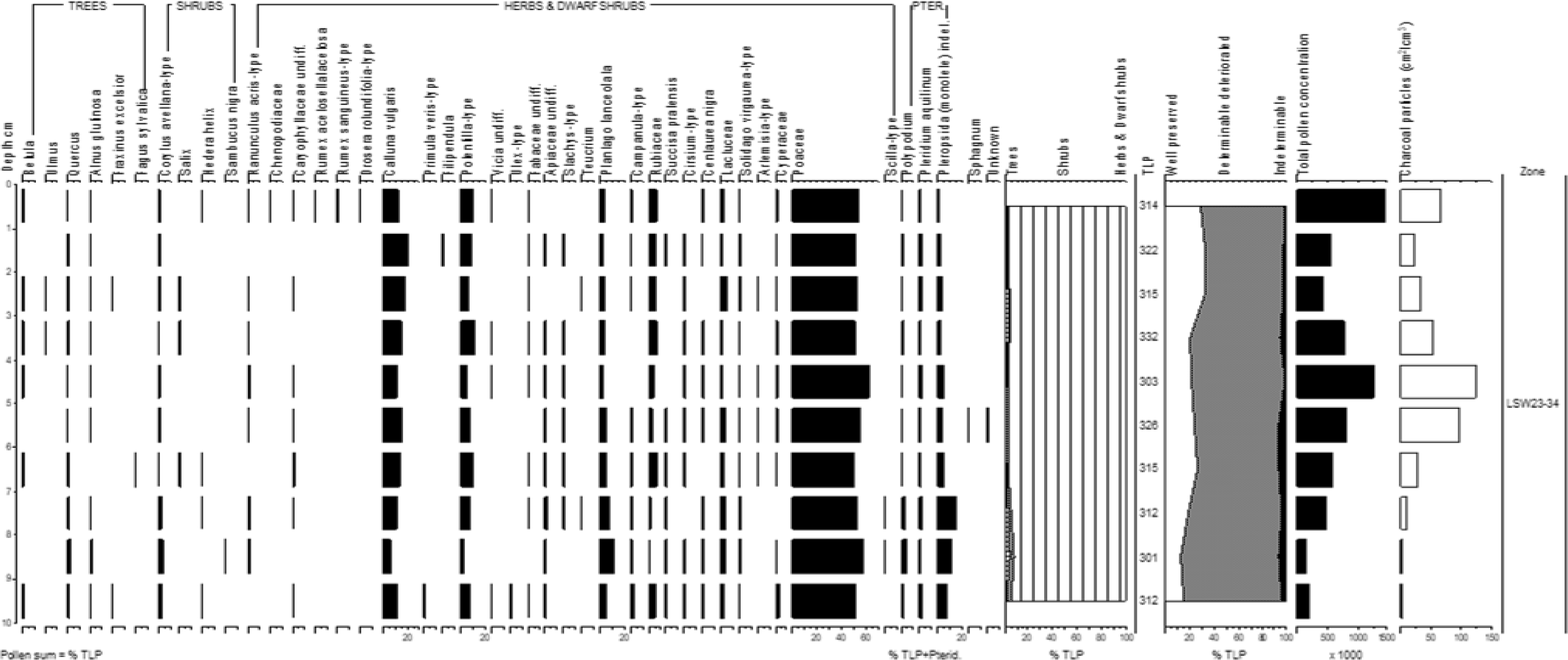
Samples from beneath stone 34 of wall of House 23 West Settlement (LSW23-34). Percentage pollen data, summary data, pollen concentrations and charcoal frequencies.

##### LSW23-34 10-0 cm (Poaceae, Calluna vulgaris, Potentilla-type, Plantago lanceolata, Rubiaceae zone)

With tree and shrub pollen comprising <10% TLP this assemblage is dominated by herbaceous pollen. Poaceae values exceed 50% TLP. There are no major changes in the percentage pollen data. The proportion of well preserved grains declines with depth. Both the pollen concentrations and charcoal values peak between 4-6 cm and in the uppermost sample (maximum 1,456,000 cm^3^ and 110 cm^2^/cm^3^, respectively).

The absence of any clear changes in pollen stratigraphy and the high pollen concentrations and charcoal values mid-zone (indicating the downward movement of particles) suggest this assemblage was produced within an active soil. However, when the soil was active is more difficult to determine. It may have been prior to the construction of the wall (and therefore open conditions are indicated), though the decline in pollen preservation with depth cautions that it could have been recently, possibly as a result of small mammal activity.

#### 3) Samples from beneath possible cobbling outside the house

Samples (0-5 cm) were collected from a silt (context 47) beneath a possibly cobbled (cobble 7) surface found outside House 23, to help determine whether the cobbling effect was natural (periglacial) or anthropogenic in origin. One LPAZ (LSW23-7) has been defined (Figure 15).

**Figure 15:**
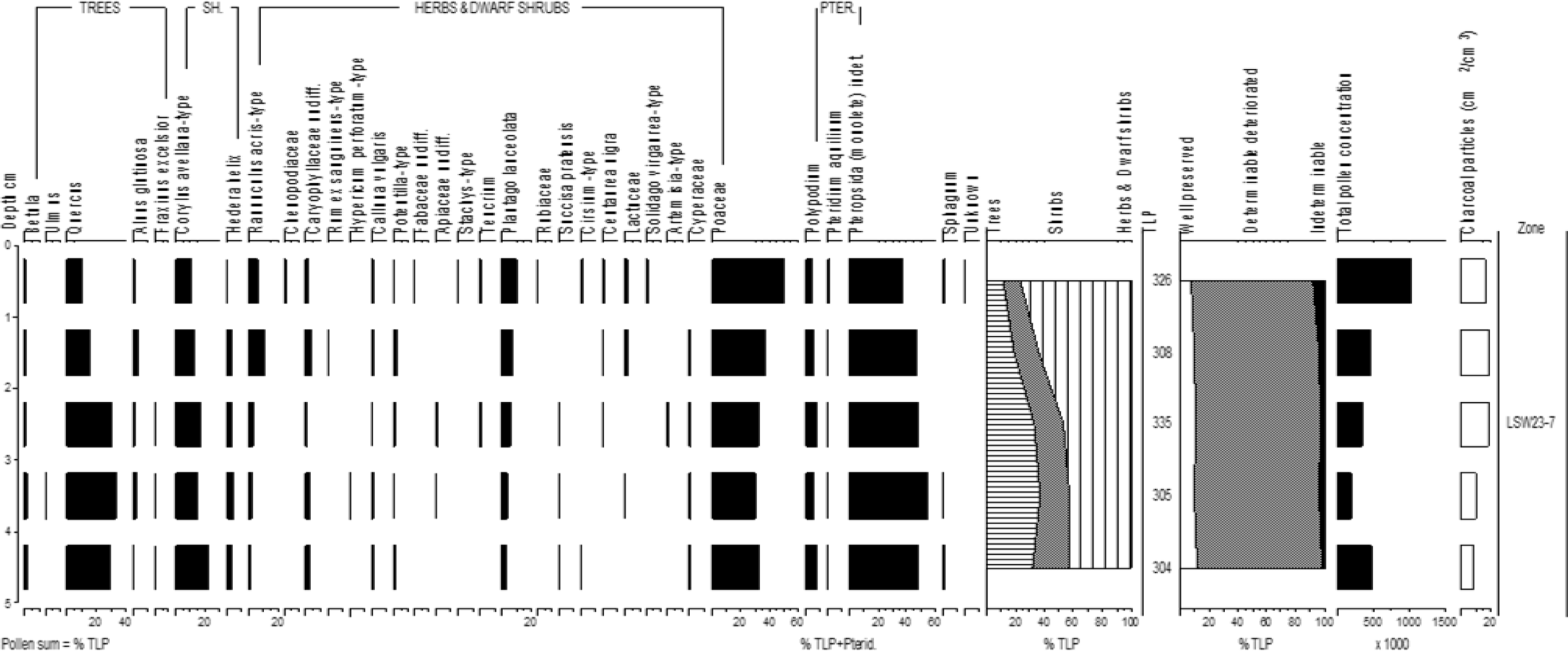
Samples from beneath possible cobbling outside House 23 West settlement (LSW23-7). Percentage pollen data, summary data, pollen concentrations and charcoal frequencies.

##### LSW23-7 5-0 cm (Poaceae, Quercus, Corylus avellana-type, Plantago lanceolata, Ranunculus acris-type zone)

Tree and shrub pollen, initially high (>50% TLP), declines above 2 cm. *Quercus* values fall from *c*. 25% to 10% TLP and *Corylus avellana*-type from 23% TLP to 10% TLP. Conversely values for Poaceae (maximum 50% TLP), *Plantago lanceolata* (maximum 11% TLP) and *Ranunculus acris*-type (maximum 10% TLP) increase at the top of the sequence. Pteropsida (monolete) spores are abundant throughout. Pollen preservation is uniformly poor (with < 12% TLP well preserved). Pollen concentrations are highest in the upper and lower-most samples and charcoal concentrations are higher in the top 3 cm.

This assemblage demonstrates that pollen has become incorporated into the silt under the ‘cobble’ during the Holocene. It resembles the assemblage from under stone 8 (LSW23-8) suggesting *Quercus* was more abundant in the area before pollen was incorporated into the upper layers from a more open landscape. If the context is sealed, then it seems likely that the cobbled surface is of anthropogenic origin and the vegetation nearby was largely open prior to its construction. The high *Ranunculus acris*-type values (which can indicate disturbance) support such an interpretation.

### Western Settlement (LSW) House 1

A large house (*c.* 12 m across) from which there is evidence for two phases of occupation (Bender *et al*. 2007). Middle Bronze Age dates were obtained from a pit and posthole and one Late Bronze Age radiocarbon date from a hearth (Table 4).

#### 1) Samples from beneath an interior wall tumble

These samples were taken from a surface buried beneath an interior wall tumble. The depths have been taken from the modern ground surface, samples from 27 and 30.5 cm are from what is thought to have been the original house floor (context 109/111) while 34 cm is from the underlying soil (context 46/109/111). One LPAZ (LSW1-44) has been defined (Figure 16).

**Figure 16:**
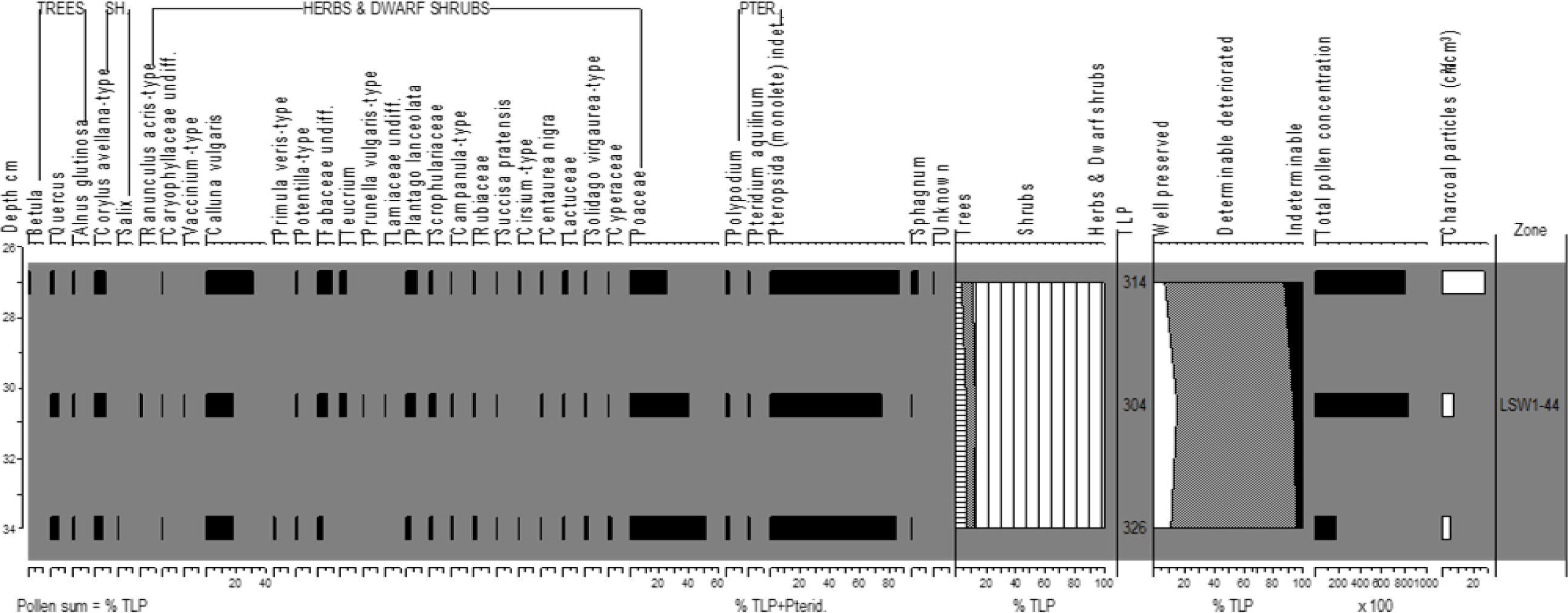
Samples from beneath interior wall rubble of House 1, West Settlement (LSW1-44). Percentage pollen data, summary data, pollen concentrations and charcoal frequencies

##### LSW1-44 34-28 cm (Poaceae, Calluna vulgaris, Pteropsida (monolete) indet. zone)

Tree and shrub frequencies are low (< 15% TLP) throughout, while values for Poaceae decline upwards (from 52% TLP to 25% TLP) and *Calluna vulgari*s increase (from 17% TLP to 31% TLP). Unusually high *Teucrium* (*c*. 4% TLP) and Fabaceae undiff. (maximum 10% TLP) values were recorded from context 46/109/111. However, this assemblage is dominated by Pteropsida (monolete) indet. spores (at *c*. 80% TLP+Pterid.). Pollen preservation is poor with <15% TLP well preserved, most of the deteriorated grains were either folded or broken. Pollen concentrations and charcoal frequencies increase upwards attaining 80,000 cm^3^ and 28 cm^2^/cm^3^, respectively, at 27 cm.

All the samples in this assemblage, following the approach of Bunting and Tipping (2000), are considered likely to have undergone post-depositional alteration and may therefore not be representative of vegetation community/communities from which they were derived. Nevertheless, the high *Calluna vulgaris*, *Teucrium* and Pterosida (monolete) indet. spores values are all consistent with abandonment. *Teucrium scorodonia* (the species from which the pollen is highly likely to have been derived) is particularly associated with scree and is sensitive to grazing (Grime *et al*. 1988). The assemblage suggests that either; the floor remained exposed with pollen becoming incorporated into the surface and to a lesser extent the underlying soil, following the abandonment of the house, or that much of the pollen has been derived from the vegetation of the wall tumble.

#### 2) Samples from wall fill and buried surface

These samples were collected from a fill between the inner and outer walls of the house (26 cm, context 15) and a soil buried (30, 34 cm and 39 cm context 49) beneath this fill. The depths given are from the modern ground surface. One LPAZ (LSW1-33) has been defined (Figure 17).

**Figure 17:**
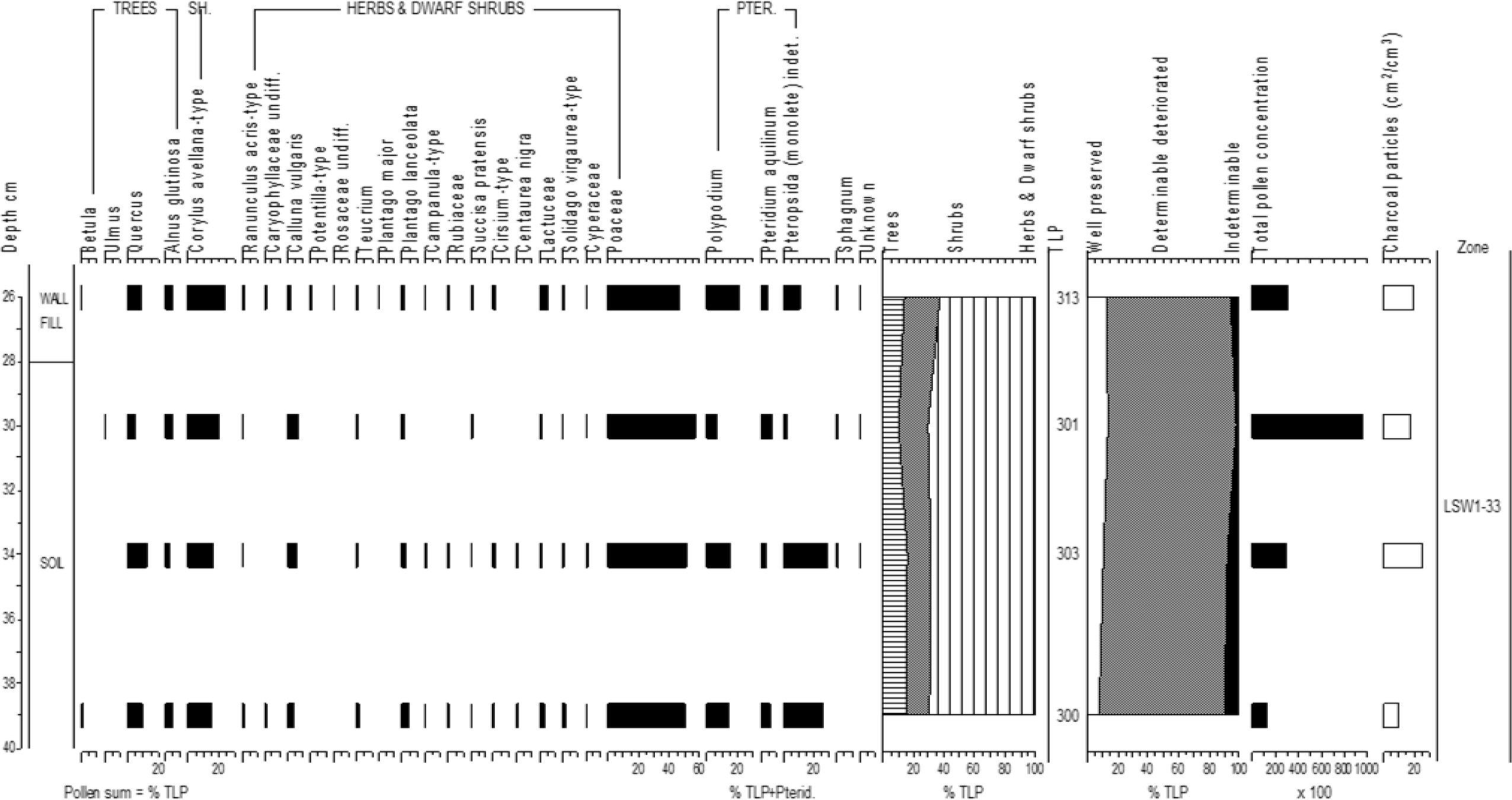
Samples from wall fill and a surface buried beneath, at House 1, West Settlement (LSW1-33). Percentage pollen data, summary data, pollen concentrations and charcoal frequencies.

##### LSW1-33 39-26 cm (Poaceae, Corylus avellana-type, Quercus zone)

Tree and shrub pollen exceeds 30% TLP, with *Corylus avellana*-type the main taxon. Poaceae values of >40% TLP are recorded throughout. Pteridophyte spores (*Polypodium* and Pteripsida (monolete) indet. are particularly well represented at 34 and 39 cm. Pollen preservation is poor with <15% TLP well preserved. Pollen concentrations peak (95,000 cm^3^) at 30 cm.

These samples form a relatively homogeneous assemblage, with the pollen concentrated in the upper levels of the buried soil. This suggests that the pollen in the soil and wall fill may have been deposited roughly contemporaneously with soil possibly being incorporated into the wall fill. The high *Corylus avellana*-type values are consistent with areas of woodland persisting, within a largely open landscape, prior to the house construction.

### Southern Settlement (LSS) House 39

A house with an internal diameter of *c.* 6.5m from which two Middle Bronze Age radiocarbon dates were obtained (Bender *et al*. 2007, Table 4).

#### 1) Samples from the interior of House 39 beneath either fallen rubble or a flagstone surface

Samples were collected from directly beneath what may have been either fallen rubble or a flagstone surface. The top samples may represent a former occupation surface (floor level), the samples beneath are from the underlying soil. Two LPAZs (LSS39-2) have been defined (Figure 18).

**Figure 18:**
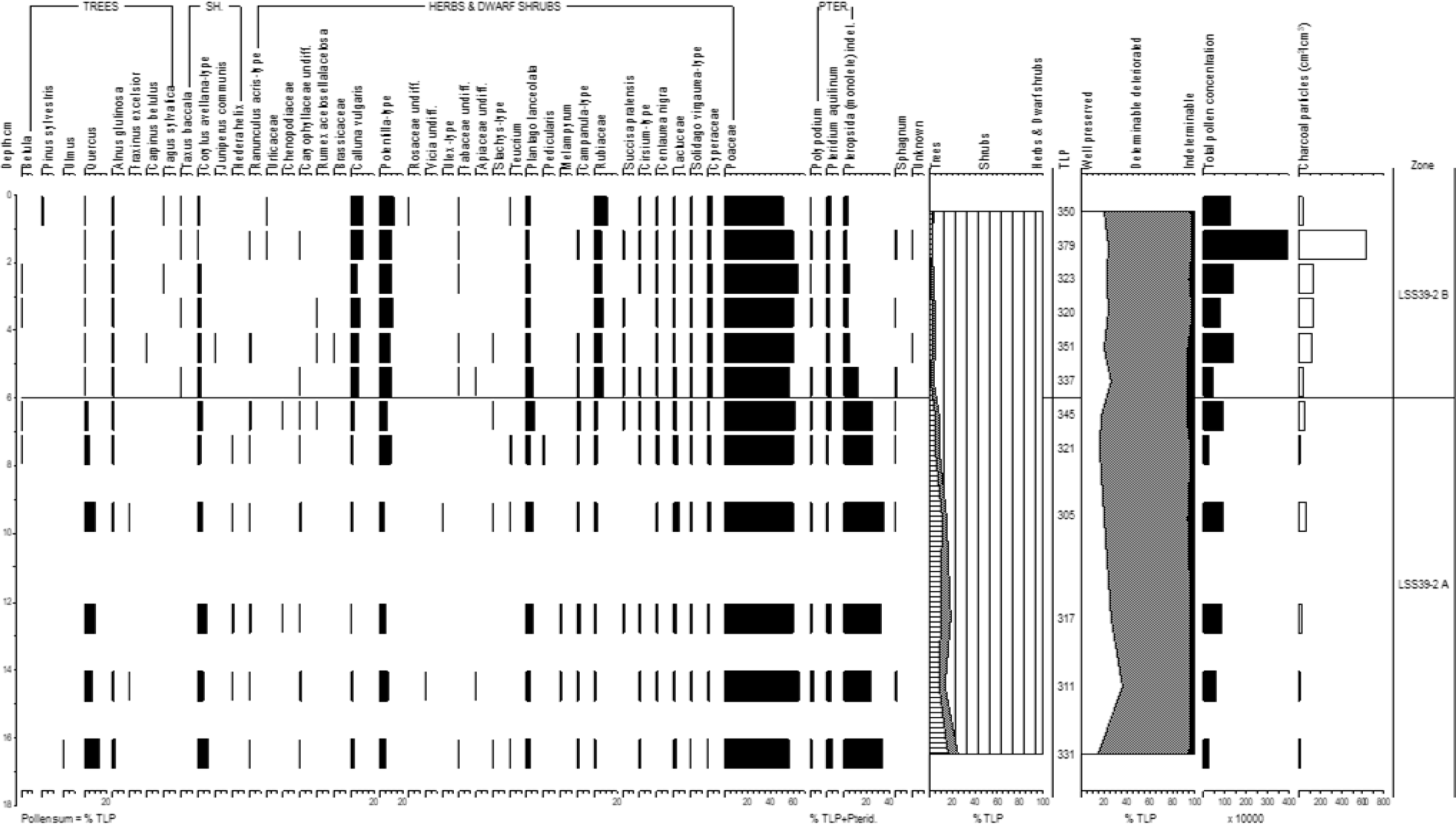
Samples from the interior of house, beneath either fallen rubble or a flagstone surface, Southern Settlement House 39 (LSS39-2). Percentage pollen data, summary data, pollen concentrations and charcoal frequencies.

##### LSS39-2 A 17-6 cm (Poaceae, Quercus, Corylus avellana-type, Pteropsida (monolete) indet. zone)

Herbaceous pollen dominates this zone (>75% TLP) with Poaceae values high throughout. *Potentilla*-type and *Plantago lanceolata* are also well represented (*c*. 5% TLP). *Quercus* and *Corylus avellana*-type pollen, at 14% TLP and 9% TLP respectively at the base, show a declining upwards trend. Pteropsida (monolete) indet. spores are common (>20 %TLP+Pterid.). Well preserved grains peak at 14.5 cm (35% TLP) but preservation is generally poor. Pollen concentrations are <1,000,000 cm^3^ and charcoal concentrations <60 cm^2^/cm^3^.

##### LSS39-2 B 6-1 cm (Poaceae, Calluna vulgaris, Potentilla-type, Rubiaceae zone)

Herbaceous pollen forms >90% TLP. Poaceae alone forms >50% TLP and *Calluna vulgaris*, *Potentilla*-type and Rubiaceae values all exceed 5% TLP. Pollen preservation improves with *c*. 25% TLP well preserved. Both pollen concentrations (3,905,000 cm^3^) and charcoal values (637 cm^2^/cm^3^) peak at 1.5 cm.

The lower LSS39-2A assemblage, containing higher tree, shrub and Pteropsida (monolete) indet. values, appears to be derived from a period when woodland cover was greater than when the upper assemblage formed. High pollen concentrations and charcoal frequencies at the top of LSS39-2B are consistent with the presence of a former land surface. The high *Calluna vulgaris*, *Potentilla*-type and Rubiaceae in LSS39-2B are indicative of heathland and acidic soil conditions and may indicate that this assemblage post-dates occupation with the overlying rubble deposited sometime after abandonment.

#### 2) Samples from a wall fill and an underlying land surface

These samples were collected from the wall fill of the double skinned house wall (4, 7 and 11 cm, context 30), the interface with an underlying land surface (13 cm, context 47) and the soil underlying this surface (15, 17, 19 cm context 55). A radiocarbon date (from *Quercus* charcoal) of 1610-1410 cal. yrs BC was obtained from the land surface (Table 4). Depths are taken from the modern ground surface above the wall fill. Two LPAZs (LSS39-11) have been defined (Figure 19).

**Figure 19:**
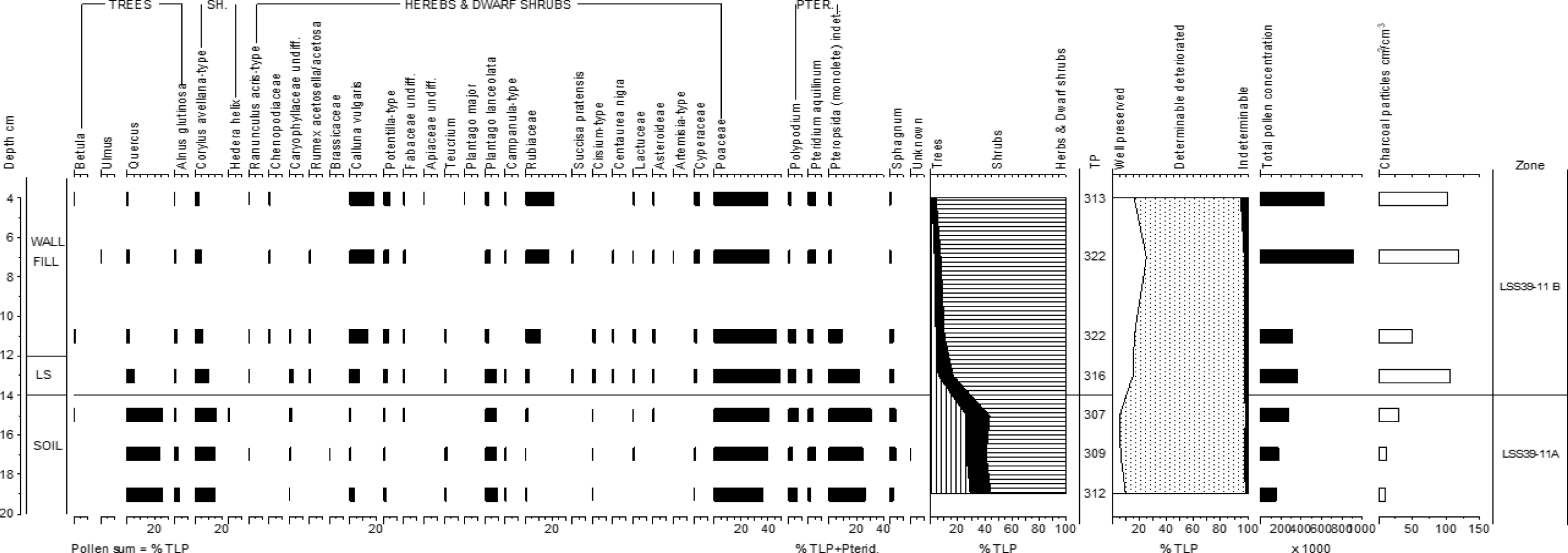
Samples from a wall fill and an underlying land surface, Southern Settlement House 39 (LSS39-11). Percentage pollen data, summary data, pollen concentrations and charcoal frequencies. LS = Land surface.

##### LSS39-11 A 19-14 cm (Poaceae, Quercus, Corylus avellana-type, Plantago lanceolata, Pteropsida (monolete) indet. zone)

Tree and shrub pollen makes up >40% TLP of the assemblage, with both *Quercus* (*c*. 25% TLP) and *Corylus avellana*-type (*c*. 15% TLP) values consistently high. Poaceae (*c*. 40% TLP) and *Plantago lanceolata* (*c*. 9% TLP) are also well represented. Pteropsida (monolete) indet. spores are common (>25% TLP+Pterid.). Pollen preservation is poor (with <12% TLP well preserved). Pollen concentrations (<160,000 cm^3^) and charcoal concentrations (<30 cm^2^/cm^3^) are consistently low.

##### LSS39-11 B 14-4 cm (Poaceae, Calluna vulgaris, Rubiaceae zone)

Herbaceous pollen makes up >80% TLP. Poaceae values, initially high, decline upwards while those for *Calluna vulgaris* and Rubiaceae rise (to maxima of 18% TLP and 21% TLP, respectively). Pollen preservation is better than in the preceding zone (with >15% TLP well preserved). Charcoal frequencies exceed 100 cm^2^/cm^3^ (except at 11 cm), while pollen concentrations are notably higher (>500,000 cm^3^) in the upper two samples.

The high tree and shrub pollen values from the soil beneath the wall (LSS39-11A) could be residual, derived from an older assemblage. However, the presence of *Quercus* charcoal dated to the Bronze Age appears to support the house being constructed while some woodland cover remained in the vicinity, with the Poaceae and *Plantago lanceolata* pollen consistent with concomitant human activity. The higher concentrations of pollen and charcoal in LSS39-11B, along with the higher values for heathland indicators, suggests the assemblage in the wall fill is not derived from the same vegetation community/communities as that which produced the pollen in the underlying soil (unlike the pollen in a similar context at House 1 Western Settlement, LSW1-33). The LSS39-11B assemblage may therefore date from after the collapse of the wall and the exposure of the fill.

#### 3) Sample from soil beneath the house wall

This (single) sample was collected from the top of the soil (1 cm context 7) beneath the house wall and should be directly comparable to the (soil) samples from context 55 above. One LPAZ (LSS39-15) has been defined (Figure 25).

##### LSS39-15 1 cm (Poaceae zone)

A zone dominated by Poaceae pollen (74% TLP). Otherwise only *Quercus*, *Corylus avellana*-type, *Calluna vulgaris* and Plantago *lanceolata* exceed 2% TLP. Well preserved grains form 13% TLP. Pollen concentrations and charcoal frequencies of 145,000 cm^3^ and 11 cm^2^/cm^3^, respectively, are recorded.

This assemblage more closely resembles that derived from context 47 (the land surface) than 55 (the soil) in the preceding investigations. If derived from a sealed context, it would suggest that the tree pollen in LSS39-11A was residual and that open conditions prevailed prior to the construction of the house wall

### Boundary Sections (BS)

The enclosure walls associated with the Leskernick settlement come in many forms, some with heaped stones and others placed with a rubble infill. Their accretional nature suggests they were constructed over a period of time. The areas enclosed range from *c*. 0.25 to 1 ha (Bender *et al*. 2007).

Buried land surfaces were widely identified in sections through the former boundaries. Generally they occur beneath an organic unit, though in places are overlain (and possibly sealed by) other sediment types (including the boundaries themselves). The boundaries examined (BSA, BSB, BSD and BSE Figure 2) may represent field walls or another form of enclosure.

#### 1) Field system controls (BSB)

Samples were collected from a site where only the overlying organic unit was present, in an attempt to characterise the pollen assemblages from this unit from an unsealed context. The samples 2-16 cm (0 cm is the modern surface) were all taken from the organic material (context 1) overlying the former land surface. One LPAZ (BSB98-2) has been defined (Figure 20).

**Figure 20:**
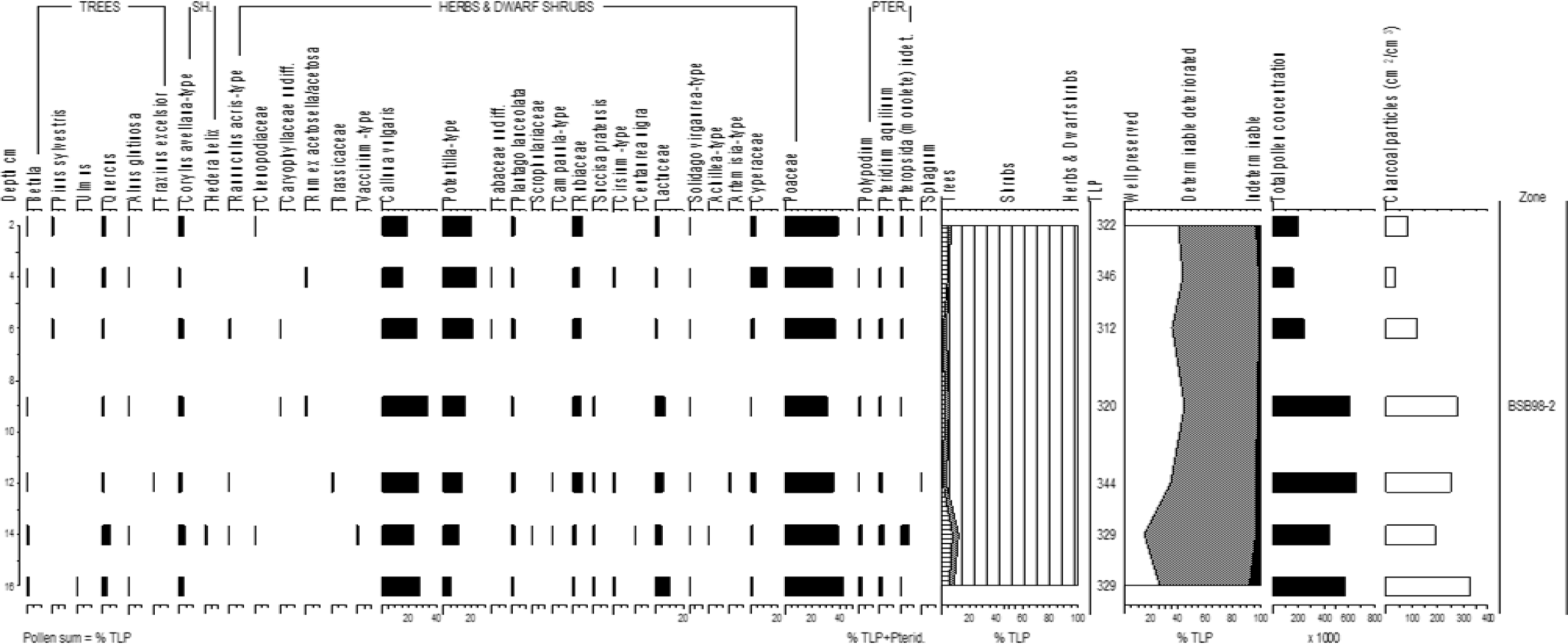
The field system control samples (BSB98-2) taken from organic sediment above the buried land surface. Percentage pollen data, summary data, pollen concentrations and charcoal frequencies.

##### BSB98-2 16-2 cm (Poaceae, Calluna vulgaris, Potentilla-type zone)

This assemblage is dominated by herbaceous pollen (>85% TLP). Poaceae values are highest (maximum 40%) at the base and top of the zone while *Calluna vulgaris* attains a maximum 31% TLP mid-zone. Frequencies for *Potentilla*-type rise from the base to 23% TLP at 4 cm. Well preserved grains form *c*. 35% TLP above 14 cm. Both pollen concentrations (maximum 652,000 cm^3^) and charcoal frequencies are notably higher below 6 cm.

The BSB98-2 assemblage is relatively homogeneous and characterised by heathland/acidic grassland indicators with both *Calluna vulgaris* and *Potentilla*-type values amongst the highest recorded from Leskernick. As would be expected in an assemblage that is receiving fresh pollen, preservation is generally good, though higher concentrations probably indicate pollen is surviving longer below 8 cm. Tree, and therefore presumably residual, pollen is notably scarce.

#### 2) Samples from the land surface and overlying wall fill (BSA)

The buried land surface was investigated where it was sealed beneath the fill of a boundary wall (between inner and outer wall stones). Samples (0 cm is the modern surface) from 15, 17, 19, 21, 24 and 27 cm are from the wall fill (context 1), 33 and 35 cm from the land surface (context 4) and 38 and 41 cm from the buried soil (context 2). Two LPAZs (BSA98-1) have been defined (Figure 21).

**Figure 21:**
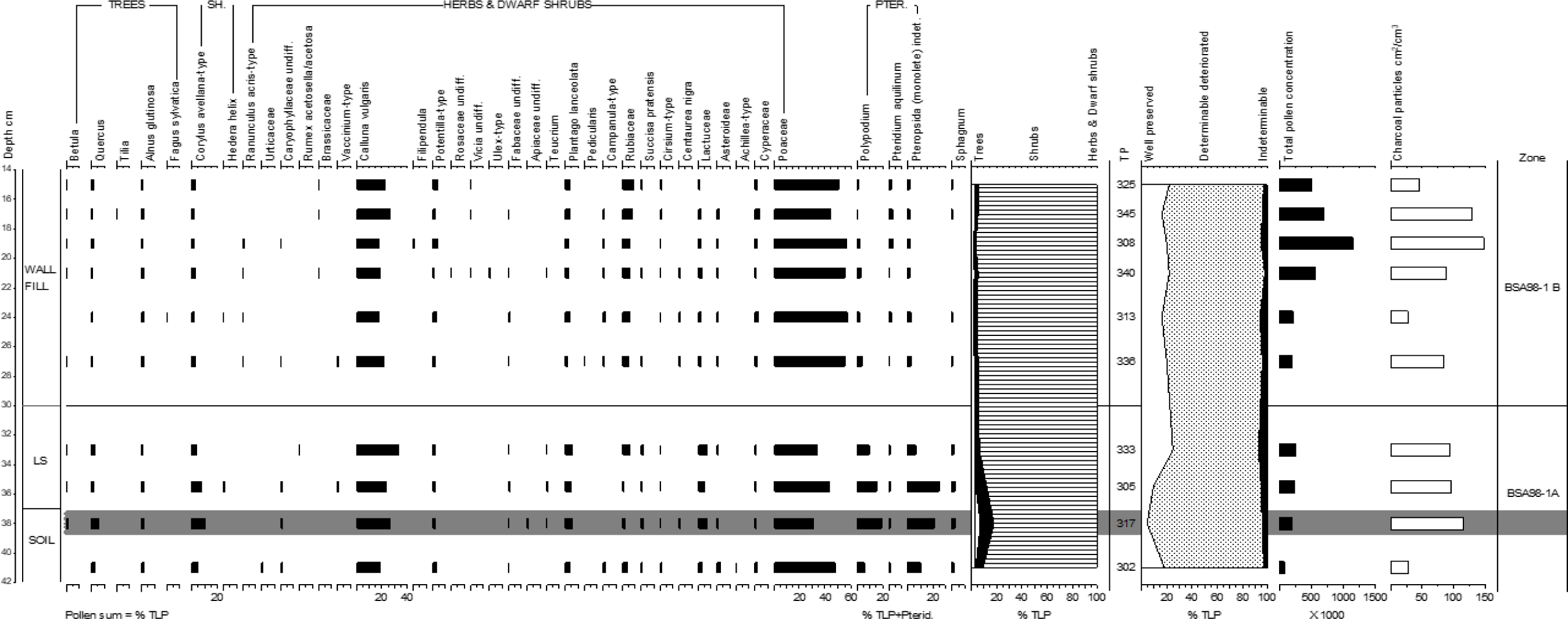
Samples from a land surface and overlying wall fill of a boundary section (BSA98-1). Percentage pollen data, summary data, pollen concentrations and charcoal frequencies. LS = Land surface.

##### BSA98-1 A 41-30 cm (Poaceae, Calluna vulgaris, Polypodium, Pteropsida (monolete) indeter. zone)

Tree and shrub pollen (largely *Corylus avellana*-type) comprises <20 % TLP. The dominant herbs are Poaceae (maximum 48% TLP) and *Calluna vulgaris* (maximum 33% TLP). Pteridophytes, both *Polypodium* and Pteropsida (monolete) indet., are well represented reaching maximum values mid-zone. The proportion of grains well preserved declines to minimum 6% TLP at 38 cm. Pollen concentrations are <210,000 cm^3^, while charcoal frequencies exceed 90 cm^2^/cm^3^ except in the basal sample.

##### BSA98-1 B 30-15 cm (Poaceae, Calluna vulgaris, Rubiaceae zone)

A more homogeneous assemblage dominated by herbaceous pollen (>90% TLP). Poaceae (maximum 57% TLP) and *Calluna vulgaris* are the main taxa, though Rubiaceae values of >5% TLP are recorded. Pteridophyte spores are scarce. Well preserved grains make up *c*. 20% TLP. Pollen concentrations rise above 24 cm (maximum 1,071,000 cm^3^) while charcoal frequencies peak (148 cm^2^/cm^3^) at 19 cm.

The assemblage from the land surface/soil is distinct with the criteria of Bunting and Tipping (2000) suggesting the sample from 38 cm may have undergone post-depositional alteration. The abundance of pteridophytes (particularly *Polypodium* which is likely to have occurred as an epiphyte on the wall/stones) and presence of *Teucrium* is consistent with much of pollen from this zone being derived from plants growing in the immediate vicinity of a boundary or clitter. The lower contexts may not then have been sealed by the construction of the wall. The wall fill assemblage is dominated by the same taxa as the control samples, though higher values for grassland taxa (Poaceae, *Plantago lanceolata*) indicate it was derived from a period of/or an area subject to more intensive grazing. Pollen concentrations rise at the top and pollen may have been incorporated into this assemblage from the modern surface.

#### 3) Samples from the land surface and overlying colluvium (BSD)

These samples were taken from a location where the land surface was buried beneath colluvium. Samples (0 cm is the modern surface) from 22 and 23.5 cm are from the colluvium (context 2), the remaining samples are from the buried soil (context 5 and 6). Two LPAZs (BSD99-4) have been defined (Figure 22).

**Figure 22:**
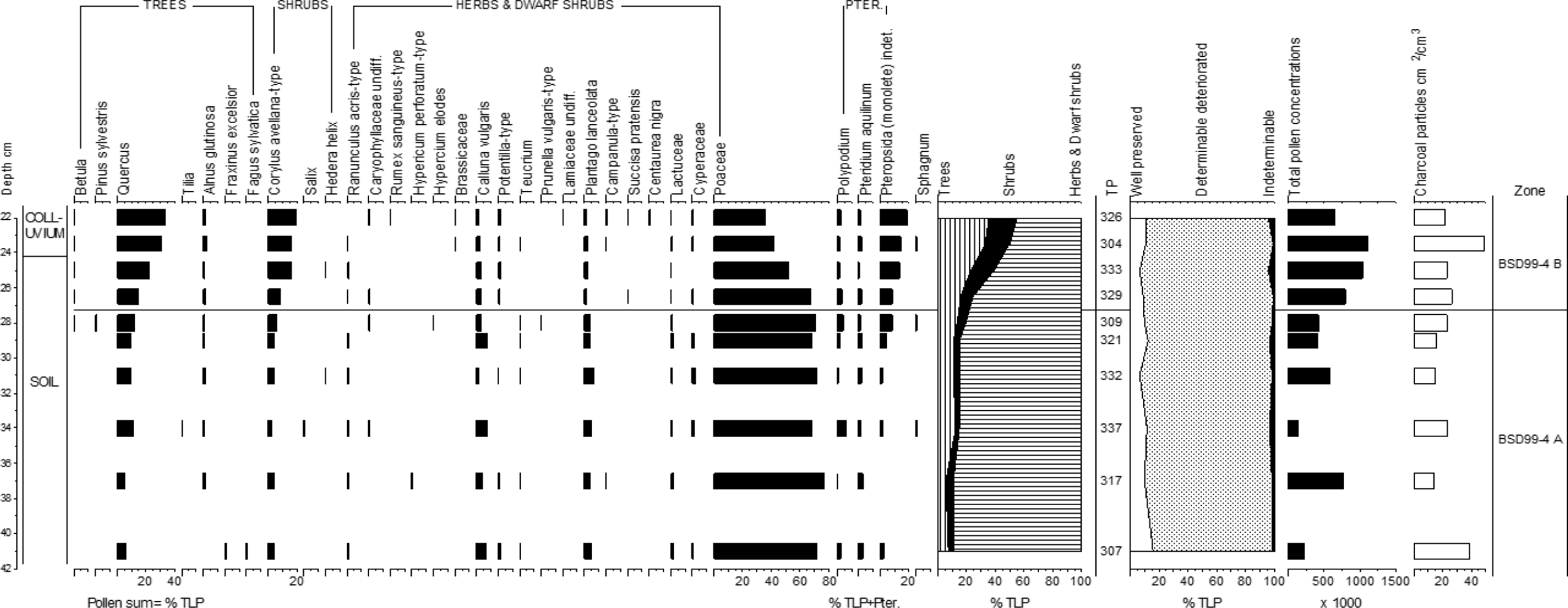
Samples from a land surface and overlying colluvium associated with a boundary section (BSD99-4). Percentage pollen data, summary data, pollen concentrations and charcoal frequencies.

##### BSD99-4A 41-27cm (Poaceae, Quercus, Calluna vulgaris, Plantago lanceolata zone)

Herbaceous pollen dominates with Poaceae values forming *c*. 70% TLP and *Calluna vulgaris* and *Plantago lanceolata* values *c*. 5% TLP. Pollen concentrations are variable. Well preserved grains form <20% TLP and charcoal frequencies are generally low.

##### BSD994B 27-22 cm (Poaceae, Quercus, Corylus avellana-type, Pteropsida (monolete) indet. zone)

Values for *Quercus*, *Corylus avellana*-type and Pteridophytes rise steadily, so that trees and shrubs comprise 56% TLP by 22 cm. The zone is also distinguished by consistently higher pollen concentrations (a maximum 906,000 cm^3^).

BSD99-4 is distinct in that the highest tree and shrub values are recorded from the top of the sequence (BSD99-4B). This coincides with the top of the soil (land surface) and the colluvium. The deposition of the colluvium (and pollen incorporation into the upper part of the soil) may therefore coincide with a period of woodland regeneration or the local growth of woody taxa, possibly in association with the boundary. Alternatively much of the tree and shrub pollen in the colluvium could be residual. Pollen concentrations are high in BSD99-4B compared those in BSD99-4A and the gradual changes between the assemblages suggests either context 5 contains some colluvium or pollen from the colluvium has been transported down the sequence (see the interpretation section). The BSD99-4A assemblage, being dominated by grassland indicators (e.g. Poaceae and *Plantago lanceolata*) is relatively distinct from the field system control assemblage (BSB98-2).

#### 4) Samples from a land surface buried beneath the spoil of a leet and a wall (BSE)

These samples were collected from a land surface at a location where a leet has been cut through a wall. The land surface is buried by spoil from the leet (Series A) and the wall itself (Series B).

Series A) At this location rubble, from when a leet was cut through the wall, covers a former land surface. Samples from 45, 48 and 52 are from a highly organic layer under the spoil (contexts 3 and 4) and the sample from 55 cm from the former land surface. One LPAZ zone (BSE99-A) has been defined (Figure 23).

**Figure 23:**
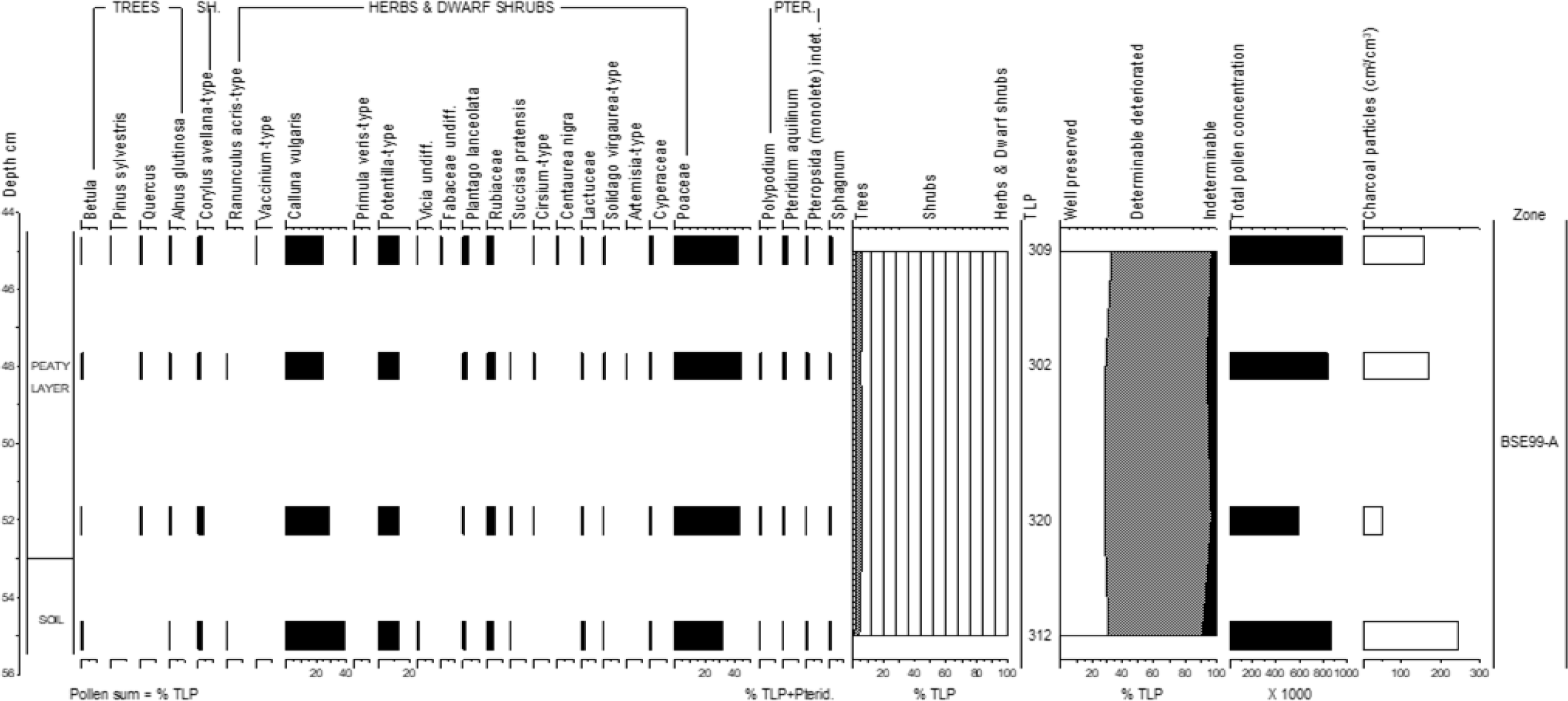
Samples from a land surface buried beneath the spoil of a leet (BSE99-A). Percentage pollen data, summary data, pollen concentrations and charcoal frequencies.

##### BSE99-A 55-45 cm (Poaceae, Calluna vulgaris, Potentilla-type zone)

An assemblage dominated by herbaceous pollen (at *c*. 95% TLP). The main taxa are Poaceae (maximum 44% TLP), *Calluna vulgaris* (maximum of 39% TLP at the base) and *Potentilla*-type (at *c*. 13% TLP). Well preserved grains make up *c*. 30% TLP. Pollen concentrations and charcoal frequencies are consistent (*c*. 850,000 cm^3^ and *c*. 175 cm^2^/cm^3^, respectively) except for a fall at 52 cm.

*Calluna vulgaris* and *Potentilla*-type values are high suggesting heathland vegetation prevailed during the formation of the organic layer. The similarity with the control assemblage (BSB98-2) is consistent with the spoil/cut being of relatively recent origin (as would be expected). The lack of a distinct assemblage from the land surface, and particularly the differences between the samples from this layer in Series A and B, indicates the downward movement of pollen into the BSE99-A sample at 55 cm may have occurred during formation of the organic layer.

Series B) The land surface sampled (at 62 cm, context 5) lies directly beneath the wall (the single sample BSE99-B is shown on Figure 25).

##### BSE99-B (Corylus avellana-type, Quercus, Poaceae, Polypodium, Pteropsida (monolete) indet. zone)

Tree and shrub pollen form 58% TLP in this sample, with *Corylus avellana*-type and *Quercus* the main taxa. Pteridophyes, both *Polypodium* (24% TLP) and Pteropsida (monolete) indet. (31%TLP), are also well represented. 8% TLP of grains are well preserved and pollen concentrations of 150,000 cm^3^ are recorded.

The land surface over which the wall was constructed contains a higher proportion of tree pollen than most of the samples collected from similar contexts (including BSA98-1 and BSB98-2). At this location the surface may therefore be older or simply less influenced by incorporation of later pollen/spores. The high *Polypodium* values may in part be a product of the poor pollen preservation, though they also suggest (if *Polypodium* was part of the wall flora) that even at this location the land surface may not have been completed sealed by the wall with some movement of pollen/spores from above.

### Boundary entrances (BE)

Samples were collected from entrances, or possibly secondary features ‘breaches’ (see Bender *et al*. 2007), in the wall boundaries at BEA and BEB (Figure 1).

#### 1) Land surface beneath entrance/breach B (BEB)

Samples were collected from a buried land surface and overlying organic layer. The samples from 7, 10 and 14 cm are from the organic layer (context 1) and 17, 20, 24 cm from the land surface (context 3). One LPAZ (BEB99) has been defined (Figure 24).

**Figure 24:**
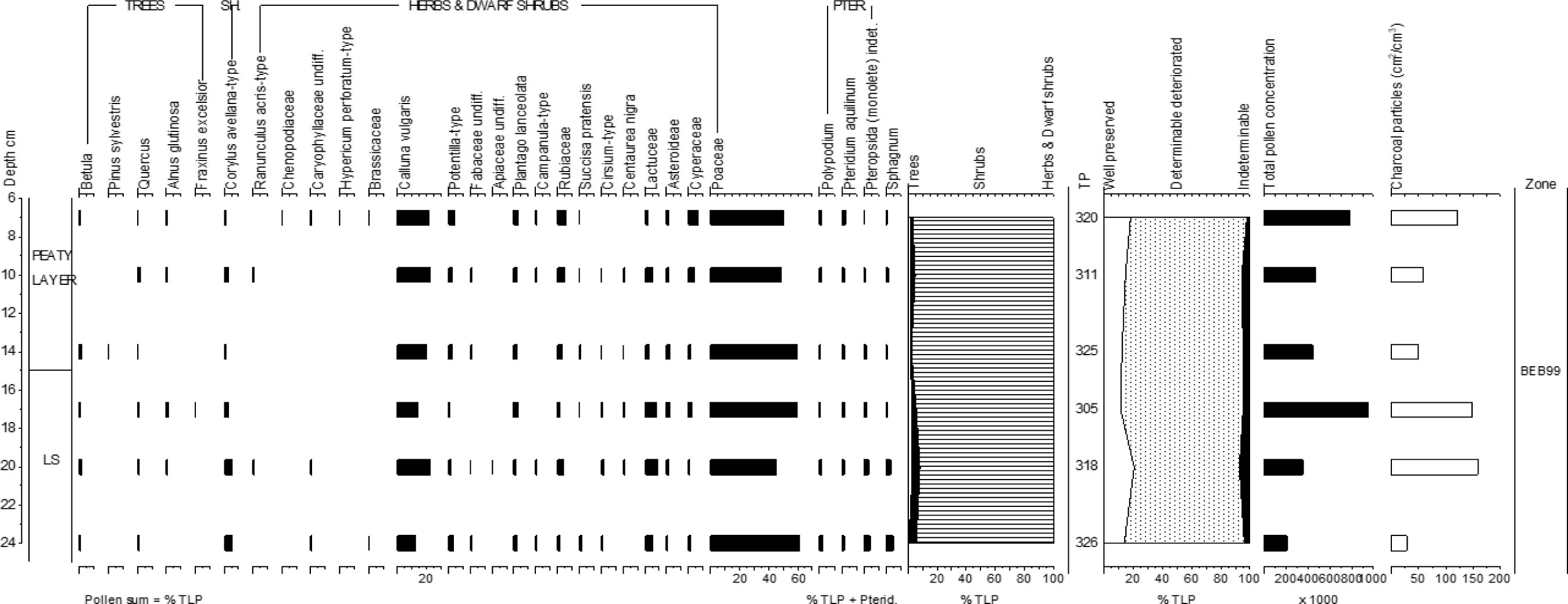
Samples from a land surface beneath a boundary entrance/breach B (BEB99). Percentage pollen data, summary data, pollen concentrations and charcoal frequencies.

##### BEB99 24-7 cm (Poaceae, Calluna vulgaris, Lactuceae zone)

A zone dominated by herbaceous pollen (>90% TLP). Poaceae (maximum 61% TLP at the base) and *Calluna vulgaris* (maximum 23% TLP at 20 cm) are the main taxa. High Lactuceae values are recorded (maximum 9% TLP) particularly below 14 cm. Over 80% TLP of the assemblage is poorly preserved. Pollen concentrations and charcoal values are highest (maximum *c*. 909,000 cm^3^ and *c*. 160 cm^2^/cm^3^, respectively) at the top and mid-zone.

High *Calluna vulgaris* pollen values from the upper, unsealed, organic layer are consistent with the field system control samples (BSB98-2) from the same unit. There is little variation in the assemblage suggesting that much of the pollen in the land surface/soil was incorporated over the same period of time as that in the peaty layer.

#### 2) Spot samples from land surface beneath entrances/breaches B (BEB) and A (BEA)

Two samples (BEB99/1 and BEB99/3) were collected from a land surface (context 3) buried under wall rubble at entrance/breach B. One sample (BEA99/9) was collected from a land surface (context 3) sealed beneath the wall at entrance/breach A. These samples are shown on Figure 25.

**Figure 25:**
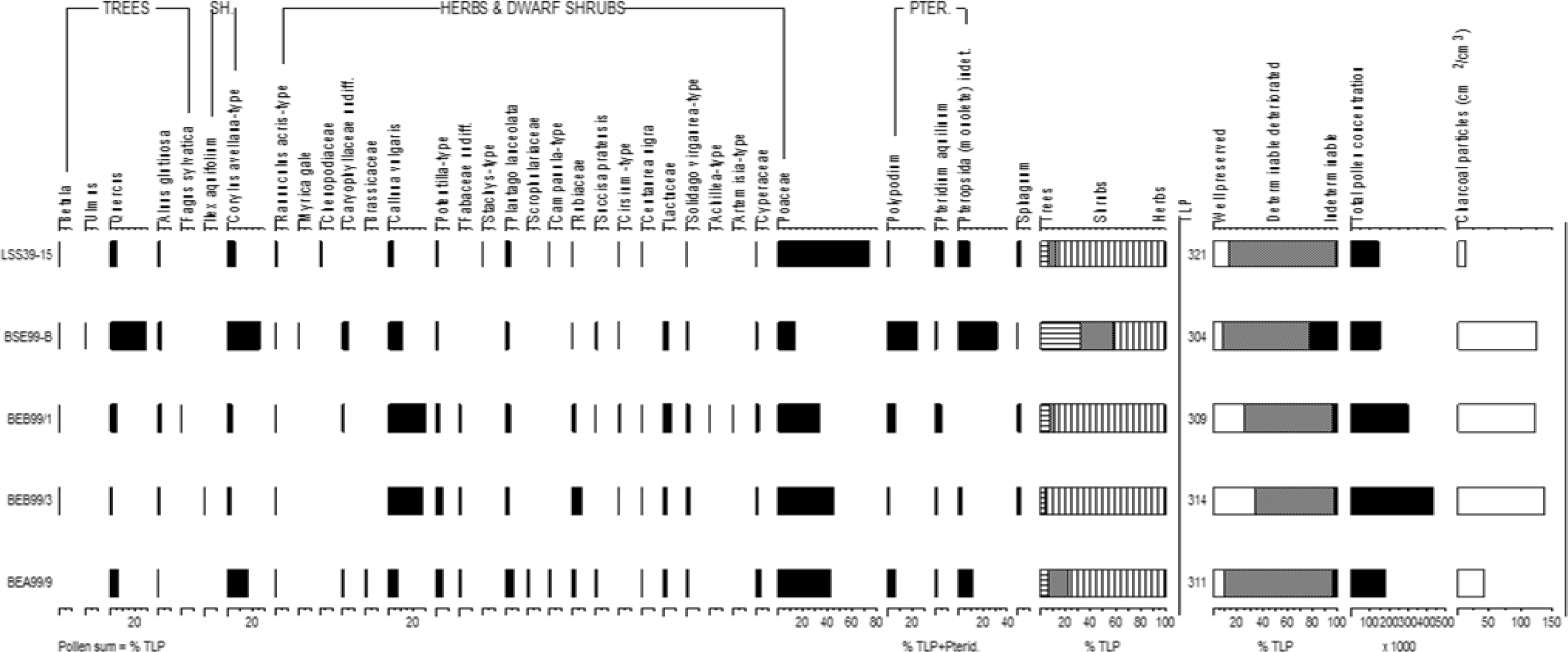
Single samples from various on-site contexts. Percentage pollen data, summary data, pollen concentrations and charcoal frequencies.

BEB99/1 and BEB99/3 assemblages are dominated by Poaceae and *Calluna vulgaris* (>25% TLP) pollen. Pollen concentrations exceed 300,000 grains cm^3^ and charcoal frequencies 150 cm^2^/cm^3^.

The spot sample from BEA99/9 also contains high Poaceae values though *Corylus avellana*-type is more abundant (16% TLP) and *Calluna vulgaris* less so, compared with the spot samples from breach B. The pollen concentration and charcoal frequencies are lower.

The samples from breach B with their high *Calluna vulgaris* values resemble the BEB99 assemblage suggesting that either the land surface was not sealed by the wall rubble or the rubble is relatively recent origin. In contrast the land surface at breach A containing more Poaceae and *Corylus avellana*-type may have been more effectively sealed beneath the wall.

### Interpretation and discussion of the on-site sequences

DCA analysis on the on-site samples was performed on all taxa that occurred at >4% TLP, excluding a number of taxa (*Hedera helix*, Caryophyllaceae, Fabaceae undiff., Scrophulariaceae, *Teucrium* and *Sphagnum*) which achieved this threshold in <5 samples. In the plot of the sample scores (Figure 26b) the context of the samples has been distinguished.

**Figure 26:**
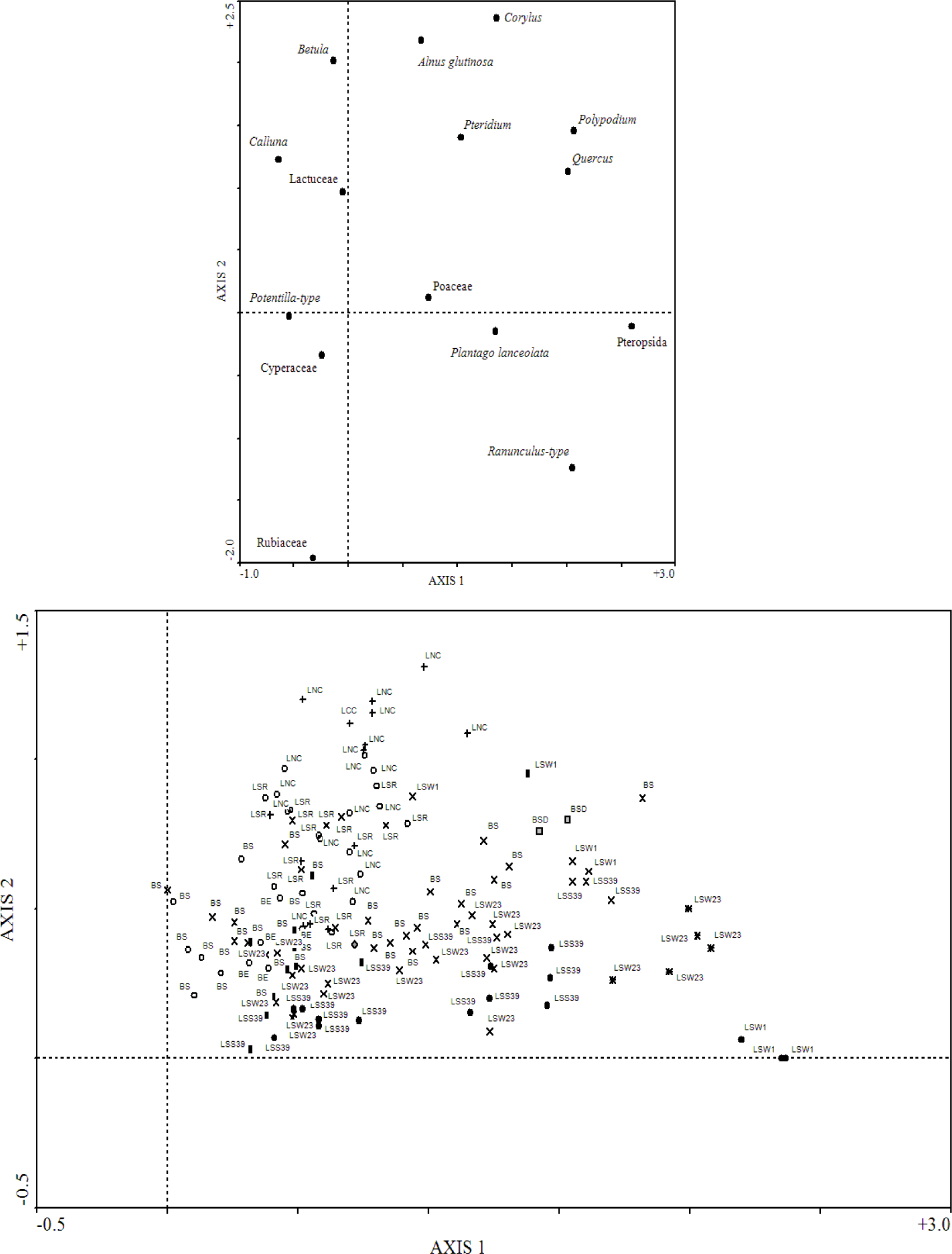
Plots of the DCA axis 1 and axis 2 scores from the on-site samples. a species scores, b site scores with the symbols indicating the sample context (post-Bronze age ‘organic’ layer and the soil beneath = ○; land surfaces and soils buried beneath archaeological/ natural features = **×**; land surfaces and soils interior to houses = •, the fill of stone holes = +, wall fill = ▪; beneath ‘cobbled’ surface = җ; from ‘spoil’ = grey diamond; from colluvium = grey square. LNC/LSR (Ritual complex) LSW (Western Settlement), LSS Southern Settlement), BS (Boundary sections), BE (Boundary entrances) BSD (Colluvium above boundary land surface)

It is difficult to interpret axis 1 in the species biplot in terms of vegetation communities (Figure 26a). However, in the samples biplot (Figure 26b) a number of the contexts are distinct. Samples with high axis 1 scores are generally derived from minerogenic material (soil beneath surfaces/colluvium), while those with low axis 1 scores are from organic layers and fills. The interpretation axis 2 is more straightforward. High axis 2 scores are associated with woodland/shrub (e.g. *Corylus avellana*-type, *Betula*) and low grazing pressure (e.g. *Pteridium aquilinum*), while the grassland and disturbance indicators (e.g. Rubiaceae, *Ranunculus acris*-type) generally have low axis 2 scores.

The samples with the highest axis 1 scores are the three samples from LSW1 (44) which are considered most likely to have been post-depositionally altered. Along with the high axis 1 values for resilient Pteropisida spores, this implies axis 1 relates to differential pollen preservation. However, the presence of taxa such as *Fraxinus*, *Taxus baccata* and the high *Quercus* pollen values at base of a number of sequences, suggests a more complex explanation, as the pollen of these taxa are particularly susceptible to decay (Havinga 1964; 1967, 1971). In addition, the criteria of Bunting and Tipping (2000) suggest, despite poor pollen preservation, the on-site samples can broadly be considered reliable.

The lowest axis 1 sample scores are derived from the boundary sections where an organic layer overlies a former land-surface. These samples contain high *Calluna vulgaris* and Cyperaceae values. Clearly originating in acidic/wet conditions, they may have accumulated pollen over considerable periods of time.

The samples derived from land surfaces/soil buried beneath archaeological/natural features occupy intermediate axis 1 positions and a number show consistent stratigraphic change. The axis 1 scores in the LSW23-8, and LSW23-7 and BSD99-4 profiles change sequentially down-profile. There is evidence to suggest both of the previously mentioned mechanisms (temporal stratification and down-washing). Higher *Quercus* pollen values with depth (e.g. down the LSW23-8 and LSW23-7 sequences) could be the result of an early phase (with mull humus) when pollen was being more actively incorporated into the sub-soil leaving a crude temporal signal. However, for the gradual decline in *Quercus* down the BSD99-4 profile, given the apparent lack of mechanisms for down-washing (Davidson *et al*. 1999; Tipping *et al*. 1999), the incorporation of colluvial sediment into the soil may be a more likely explanation.

The LNC samples from the stone-hole fill are the only on-site samples from a secure Early Bronze Age context and with high axis 2 scores they contain the highest proportions of tree/scrub pollen. In general the high axis 2 scores from the ritual complex (LNC and LSR) suggest, along with the LESKM-1 assemblage, that woody taxa were more abundant in the Late Neolithic/Early Bronze Age than during subsequent occupation phases and that the early abandonment of the complex was followed by a period of low grazing pressure.

The samples from interior of the houses are distinguished by low axis 2 scores, with high percentages of Poaceae, *Plantago lanceolata* and *Ranunculus acris*-type. This can be interpreted as the sediments sampled incorporating a high proportion of their pollen content during a period of relatively intense land use, presumably immediately before or during their occupation.

The assemblages from the boundary sections of field systems might also be expected to indicate intense land-use, however, only the buried soil beneath colluvium (BSD99-4A) conforms to this expectation. They generally occupy intermediate axis 2 scores with high *Calluna vulgaris* values. In some cases this may reflect conditions prior to wall construction indicating the heavily clittered southern slope of Leskernick was relatively under-grazed. Alternatively, given the potential for the destruction and reuse of the enclosure walls, these assemblages may have formed during or after the extended period over which the settlement was abandoned (from the later Middle Bronze Age onwards, Bender et al. 2007).

It should be noted that no pollen grains consistent with cereals were found in any of the on-site samples. This certainly argues against the settlement enclosures being used for cereal production.

The ‘control’ samples were collected to determine the extent to which contexts of archaeological interest (the Stone Row and Boundary Sections) may have become contaminated by more recently deposited pollen. These assemblages are not distinct in the DCA. Part of the difficultly in this approach (and interpreting the Leskernick assemblages in general) is that moorland/grassland communities appear to have present, to a greater or less extent, in the Leskernick area since the late Neolithic. In many, the dominant taxa cannot be palynologically distinguished to species level. Recently introduced ‘exotic’ taxa (e.g. conifers) are locally absent. Also of concern is that differences between the control assemblages and those from the contexts of interest may reflect local-scale spatial variations in vegetation rather than temporal differences. Work on pollen-vegetation relationships in a moorland setting suggests that much of the pollen from non-arboreal taxa (including *Calluna vulgaris*, Poaceae, *Potentilla*-type) derives from an area within a radius of 2 m or less (Bunting 2003). In the heterogeneous clittered landscape of the settlement and enclosures, differences between on-site assemblages, even those obtained in relatively close proximity, could simply be the result of spatial variations in a broadly contemporary vegetation mosaic.

## Summary of vegetation/land-use changes and Conclusion

Figures 27 and 28 summarise the vegetation/land use changes inferred from the pollen sequences. It is possible to create a coherent narrative for the Leskernick area which is consistent with the archaeological evidence (Figure 27) though to a lesser extent with the historical evidence for the wider area (Figure 28). The chronological uncertainties are a major constraint, alongside the difficulties in interpreting the minerogenic on-site samples. That no continuous sequence spanning the Neolithic to the present was discovered is disappointing, but unsurprising given the length of time, and the nature and degree to which the Leskernick landscape and Bodmin Moor in general have been exploited.

**Figure 27:**
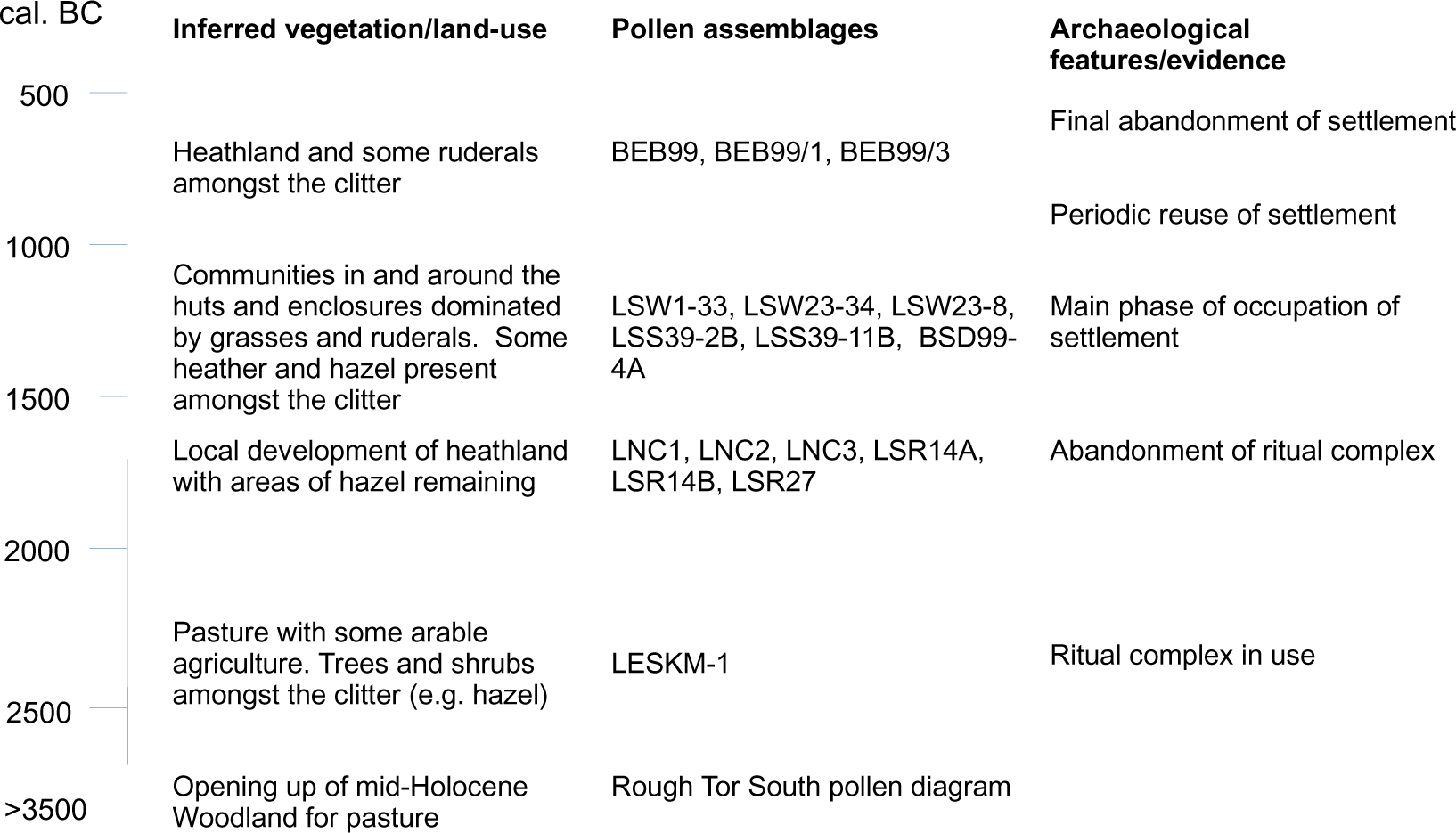
The vegetation inferred from the pollen evidence against the archaeological evidence for the prehistoric period.

**Figure 28:**
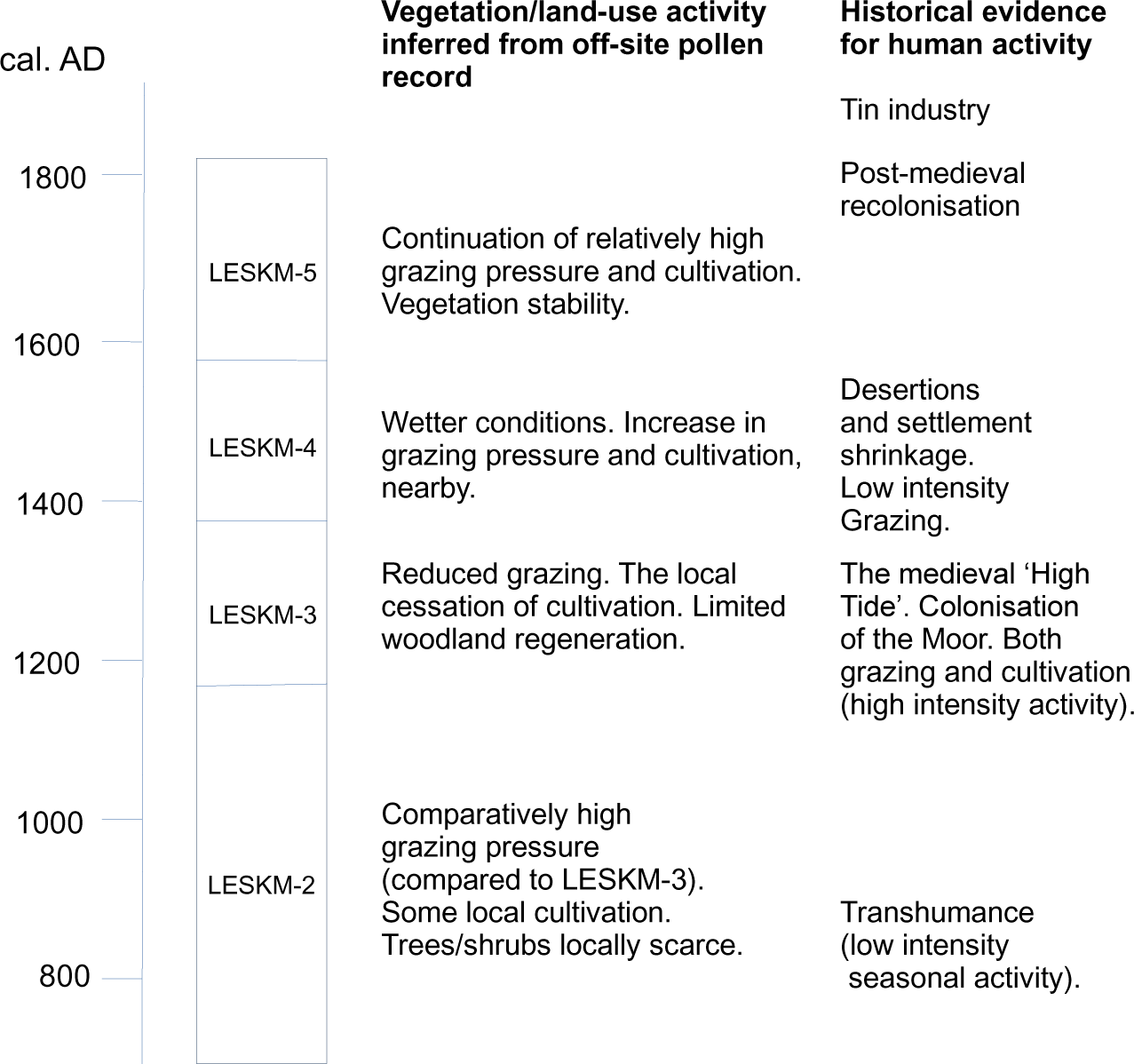
Vegetation/Land use history inferred from the Leskernick off-site pollen record compared against the historical evidence for Bodmin Moor.

## Acknowledgements

The on-site samples were provided by the excavation team led by Sue Hamilton. The opinions expressed are those of the author alone.

